# Specialized gas-exchange endothelium of the zebrafish gill

**DOI:** 10.64898/2025.11.30.690480

**Authors:** Jong S. Park, Justin Gutkowski, Daniel Castranova, Madeleine Kenton, Abhinav Sur, Celia Martinez-Aceves, Minh-Anh Nguyen, Christopher Dell, Gennady Margolin, James Iben, Louis Dye, Melissa R. Mikolai, Adam Harned, Van Pham, Ryan K. Dale, Jeffrey A. Farrell, Kedar Narayan, Brant M. Weinstein

**Affiliations:** Division of Developmental Biology, National Institute of Child Health and Human Development, National Institutes of Health, Bethesda, MD, 20892; Bioinformatics and Scientific Programming Core, Eunice Kennedy Shriver National Institute of Child Health and Human Development, NIH, Bethesda, MD 20814, USA; Molecular Genomics Core, Eunice Kennedy Shriver National Institute of Child Health and Human Development, NIH, Bethesda, MD 20814, USA; Microscopy and Imaging Core, Eunice Kennedy Shriver National Institute of Child Health and Human Development, NIH, Bethesda, MD 20814, USA; Center for Volume Electron Microscopy, Center for Cancer Research, National Cancer Institute, National Institutes of Health, Bethesda, MD 20892, USA; Cancer Research Technology Program, Frederick National Laboratory for Cancer Research, Frederick, MD 21702, USA

**Keywords:** Lung, Gill, Gas-exchange blood vasculature, Oxygen-exchange, Zebrafish, Aerocytes, Cap2, General capillary, Cap1

## Abstract

The pulmonary vasculature plays a critical role in gas exchange and in lung pathologies, but it is challenging to observe and experimentally manipulate deep within the lungs of living mammals. Unlike mammalian lungs, externally located zebrafish gills are readily accessible for high-resolution optical imaging and experimental manipulation, suggesting zebrafish might provide an excellent comparative vertebrate model for studying the development and function of gas-exchange organs and the gas-exchange blood vasculature. To characterize their resident cell populations, we performed single-cell RNA sequencing (scRNAseq) on adult zebrafish gills, revealing numerous cell types with transcriptional similarities to those found in mammalian lungs. We uncovered and characterized several different endothelial cell populations, including distinct clusters of arterial endothelial cells and lymphatic endothelial cells. The largest endothelial cell cluster closely resembles Cap2 or “Aerocyte” endothelial cells, a recently discovered unusual mammalian endothelial cell type found exclusively in lung alveoli. Zebrafish aerocytes localize to the analogous gas-exchange structures in fish, the highly vascularized gill lamellae. We use confocal and super-resolution imaging of transgenic and hybridization chain reaction-probed zebrafish, array tomography, and focused ion beam scanning electron microscopy to carry out a detailed and comprehensive characterization of gill aerocytes including 3-D ultrastructural reconstruction of one of these cells, showing that as in mammals these cells are closely associated with gas-exchange epithelia and that they possess unique properties that may help facilitate their gas-exchange function. Together, our findings help establish a new, experimentally accessible comparative vertebrate model for studying the gas-exchange blood vasculature.

## INTRODUCTION

Vertebrates have developed specialized organs for gas exchange to facilitate the uptake of oxygen and the removal of carbon dioxide. In mammals, lungs and lung alveoli provide an enormous surface area for diffusion of oxygen and carbon dioxide across the thin blood-air barrier. The vasculature is crucial for gas exchange and transport to and from the lungs. Vascular endothelial cells are one of the most abundant cell populations in the lung, and emerging evidence suggests they play a central role in many different vascular pathologies, including acute respiratory distress syndrome (ARDS) (Huppert et al., 2019; Vassiliou et al., 2020), chronic obstructive pulmonary disease (COPD) (Screm et al., 2024; Theodorakopoulou et al., 2021), influenza (Armstrong et al., 2013; Marchenko & Zhilinskaya, 2024; Niethamer et al., 2025), and COVID-19 (Libby & Lüscher, 2020; Nappi & Avtaar Singh, 2022).

Recent single-cell RNA sequencing studies have shown that lung alveoli possess highly specialized, unusual endothelial cells named Cap2 cells or “Aerocytes,” (Ellis et al., 2020; Gillich et al., 2020; Niethamer et al., 2020). Aerocytes have a large and complex surface area and associate very closely with alveolar epithelial cells, suggesting they are critical for gas exchange function (Ellis et al., 2020; Gillich et al., 2020; Niethamer et al., 2020). However, their location deep within mammalian lungs and relative inaccessibility to live imaging and experimental manipulation make it challenging to study the development and function of these specialized gas-exchange endothelial cells in their endogenous *in vivo* context.

Similar to lungs, gills function as gas-exchange organs in vertebrates. Although gills operate in water, they share many features with lungs. Both organs perform gas exchange by diffusion within their primary gas-exchange units. In lungs, this unit is the alveolus, while in gills, the corresponding structure is called the lamella (Evans et al., 2005; Molnar & Gair, 2022; Seadler et al., 2023)(**Fig. 1A**). Gills and lungs share many cell types, including respiratory pavement epithelial cells, oxygen-sensing neuroepithelial cells, and chemosensory tuft cells (Pan et al., 2022; Smith et al., 2018). Gills and lungs also both feature extensive, highly branched vascular networks dedicated to efficient gas exchange. However, unlike lungs, which are located deep inside the bodies of mammals and other air-breathing vertebrates, gills are externally positioned and easily accessible for live microscopy and experimental manipulation in many vertebrates, including zebrafish.

**Figure 1.**
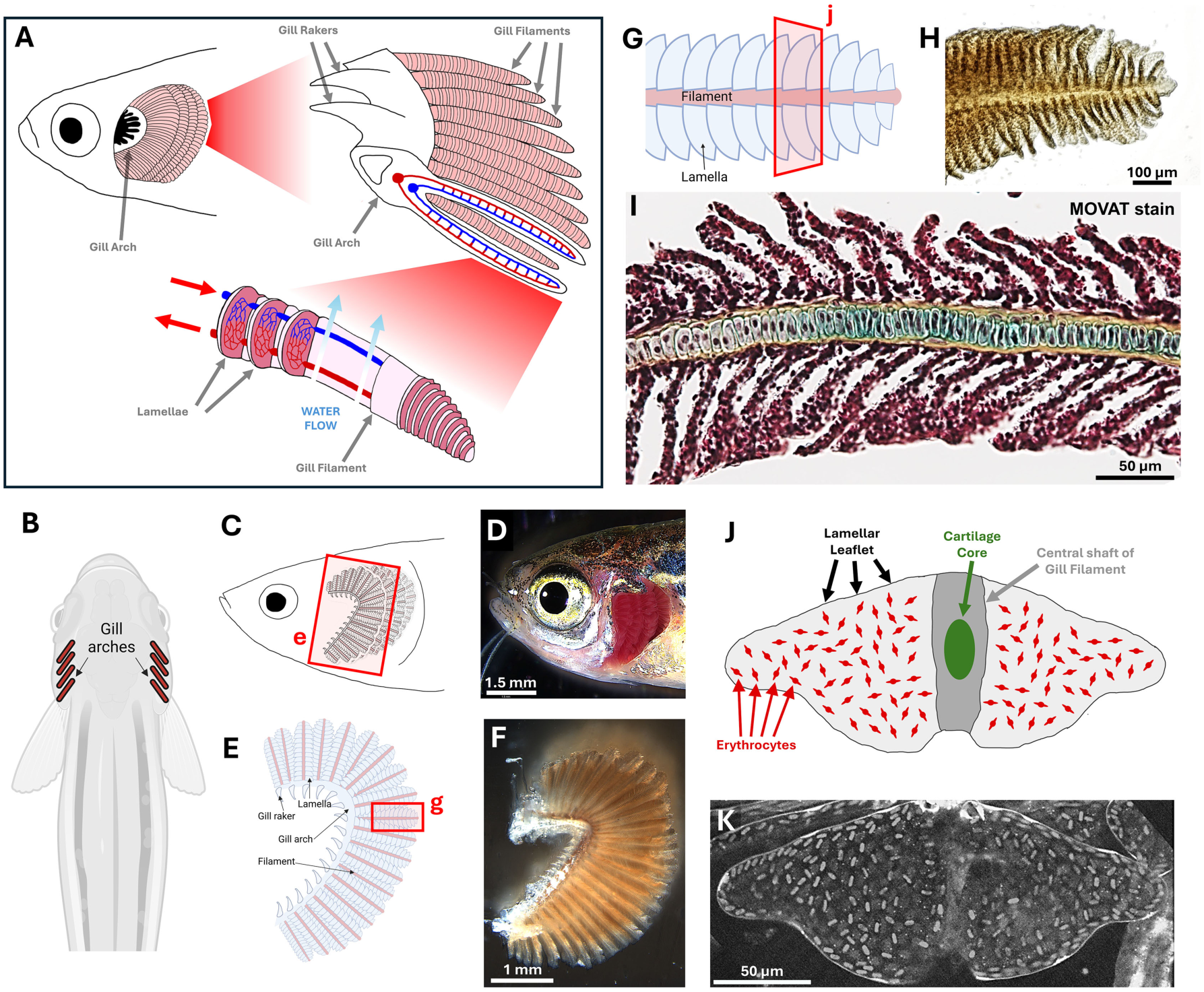
Anatomy of the zebrafish gill. **A,** Schematic diagram of adult zebrafish gill anatomy. **B,** Dorsal view schematic diagram of an adult zebrafish showing the positions of the eight gill arches, four on each side. **C,D,** Schematic diagram (C) and corresponding brightfield micrograph (D) showing a lateral view of an adult zebrafish head with operculum curled back to show gills. Box in panel C shows area depicted in panel E. **E,F,** Schematic diagram (E) and brightfield micrograph (F) of a dissected zebrafish gill arch. Box in panel E shows area depicted in panel G. **G,H,** Schematic diagram (G) and brightfield micrograph (H) of a single dissected zebrafish gill filament. Box in panel G shows area depicted in panel J. **I,** Brightfield micrograph of a sectioned zebrafish gill filament with MOVAT staining. **J,K,** Schematic diagram (J) and live confocal micrograph (K) of the zebrafish gill lamella. Scale bars = 1.5 mm (D), 1 mm (F), 100 µm (H), and 50 µm (I,K).

Zebrafish are a well-established vertebrate model organism due to their external development, genetic and experimental accessibility, optical clarity, and suitability for developmental and regenerative studies (Teame et al., 2019). Like other teleost fish, zebrafish have highly organized, complex gills whose superficial location makes them highly accessible for live optical imaging and experimentation. Earlier studies on the gas-exchange vasculature of teleost fish identified “pillar cells” as key cellular components of the gill lamellae (Evans et al., 2005; Wilson & Laurent, 2002) while more recent work in zebrafish has advanced our understanding of gill vascular patterning (Preußner et al., 2025) and linked pillar cell development to cranial neural crest-derived lineages (Fabian et al., 2022). While these studies provide important developmental insights, the precise molecular identities, cellular structure, and functional roles of the specialized cells in the gills remain to be fully elucidated.

In this study, we conduct a detailed anatomical, cellular, and molecular characterization of the zebrafish gills and their resident cell populations, with a particular focus on their endothelial cells. We utilize single-cell RNA sequencing, high-resolution imaging, histology, and volume electron microscopic ultrastructural analysis (array tomography, FIB-SEM) to characterize gill morphology and gill cell types. We report a variety of cell types resembling those commonly found in mammalian lungs, including gas exchange epithelial cells, ionocytes, goblet cells, and chemosensory cells. We also provide new information on the vascular morphology of the gills and characterize several molecularly heterogeneous endothelial cell populations and their localization within the gills. Importantly, we demonstrate that the major endothelial cell type in the gills is a lamellar endothelial cell population that resembles mammalian lung Cap2 or “Aerocyte” cells, and we describe the highly unusual features of these gas-exchange endothelial cells.

## MATERIALS AND METHODS

### Fish husbandry and strains

Fish were housed in a large zebrafish-dedicated recirculating aquaculture facility (4 separate 22,000 L systems) in 6 L and 1.8 L tanks. Fry were fed rotifers, and adults were fed Gemma Micro 300 (Skretting) once per day. Water quality parameters were routinely measured, and appropriate measures were taken to maintain water quality stability (water quality data available upon request). The following transgenic fish lines were used for this study: *Tg(mrc1a:eGFP)*^y251^ (Jung et al., 2017)*, Tg(kdrl:mcherry)^y206^* (Fujita et al., 2011)*, Tg(fli1a:eGFP)^y1^* (Lawson & Weinstein, 2002)*, Tg(5.2lyve1b:dsred)^nz101^* (Okuda et al., 2012), *Tg(flt4:yfp)^hu4881^* (Hogan et al., 2009), *Tg(gng13a:egfp)^y709^* (Castranova et al., 2025), *Tg(fli1a:lifeact)^mu240^* (Hamm et al., 2016). Some transgenic lines were maintained in a *casper* (*mitfa^w2/w2^, mpv17 ^a9/19^* double mutant) background (White et al., 2008) to enhance the visualization of the gill by eliminating melanocyte and iridophore cell populations. This study was conducted in an AAALAC-accredited facility under an active research project overseen by the NICHD ACUC, Animal Study Proposal # 24-015.

### Live imaging of the adult zebrafish gill

Adult zebrafish with defective operculum (gill cover) were anesthetized with buffered tricaine (126 mg/L), placed in an imaging dish (Lab-TekII #155360), and gently covered with a sponge to prevent movement. For longer imaging sessions, fish were intubated in a 3D-printed chamber holding an imaging dish, with tricaine water pumped directly into the fish’s mouth using a peristaltic pump at 15 rotations per minute, as described in (Castranova et al., 2022).

### Dissection of adult zebrafish gill for imaging

Fish were euthanized by submersion in an ice bath. The gill basket was removed with a pair of Dumont L5 forceps and surgical scissors. Gill arches were further separated and then dipped in 1x PBS to remove debris. For imaging, gill tissues were placed in a 35-mm glass-bottom petri dish (MatTek #P35-1.5-14-C) containing 1x PBS, with a circular coverslip placed on top.

### Light Microscopy

Approximately 3 fish were imaged per experiment. All images were analyzed, and those selected for presentation represented the complete set collected. Both male and female fish were chosen randomly for analysis.

#### Stereo Microscopy (Leica M205)

Brightfield images of the zebrafish head and gill were acquired using a Leica M205 stereo microscope and LAS V4.7 or LAS X software, with extended depth of focus z-stack image processing.

#### Confocal Microscopy (Nikon W1)

Confocal images of the gill were acquired using a Nikon Ti2 inverted microscope equipped with a Yokogawa CSU-W1 spinning disk confocal and a Hamamatsu Orca Flash 4 v3 camera. The following Nikon objectives were utilized: 4X Air, 0.2 N.A.; 10X Air, 0.45 N.A.; and 40X water immersion, 1.15 N.A. The large size of the adult zebrafish head and gill often necessitated tile acquisitions that were later stitched together using Nikon Elements software.

#### Stimulated emission depletion (STED) Microscopy

STED images of the gill were acquired using Abberior MIRAVA Polyscope. All images were taken with a 60x silicone objective and analyzed with LiGHTBOX software (v2024.48.21878).

### Transmission Electron Microscopy

Zebrafish gills were fixed in 4% glutaraldehyde prepared in 0.1 M sodium cacodylate buffer, pH 7.4. After fixation samples were inserted into mPrep tissue capsules and loaded onto an mPrep ASP-2000 Automated Biological Specimen Preparation Processor (Microscopy Innovations, LLC, Marshfield, WI.) which automates all processing steps which included the following steps: 0.1M sodium cacodylate buffer rinses, post-fixation in 2% osmium tetroxide (made in 0.1M sodium cacodylate buffer), dehydrated in a graded ethanol series followed by further dehydration in 100% acetone and finally infiltrated and embedded in Embed 812 epoxy resin (Electron Microscopy Sciences Hatfield, PA.). Embedded samples were polymerized in an oven set at 60 °C. Samples were then ultra-thin sectioned (90 nm) on a Leica EM UC7 Ultramicrotome. Thin sections were picked up and placed on 200 mesh copper grids and post-stained with UranyLess (Uranyl Acetate substitute, Electron Microscopy Sciences, Hatfield, PA.) and lead citrate. Imaging was performed on a JEOL-1400 Transmission Electron Microscope operating at 80kV and an AMT BioSprint-29 (29 megapixels) camera.

### Array Tomography and FIB-SEM

#### Sample preparation

Zebrafish were euthanized, and their gills were dissected to isolate individual gill arches. Gill arches were fixed with Karnovsky’s fixative (4% PFA + 2% GA in 0.1M sodium cacodylate buffer) and were post-fixed in 2% OsO_4_ + 1.5% potassium ferricyanide in 0.1 M sodium cacodylate for 1 hour at room temperature (RT). Next, gills were washed with ultrapure water and stained with 1% aqueous uranyl acetate for 1 hour at RT. After washing (5 repeats of 3-minute incubations) with water, the gills were treated with lead aspartate at 60°C for 30 minutes and washed again with water. The gills were dehydrated through a graded ethanol series (10 minutes each at 35%, 50%, 70%, and 95%, with 3 repeats of 10 minutes at 100%). Gills were further dehydrated in 100% propylene oxide (PO) for 3 exchanges of 10 minutes each. After dehydration, the gill arches were infiltrated with increasing amounts of Polybed resin diluted in PO (resin:PO sequential ratios of 1:3, 1:1, 3:1, and 100%). After 100% infiltration, gill arches were cut into three portions (“base”, “middle”, “top”) and placed on an aclar sheet. The Aclar sheet itself was placed within a gene frame (Thermo Fisher) on a microscope slide, and a drop of 100% degassed resin was added. Another piece of Aclar was placed atop the gene frame, essentially creating a flat, thin resin sandwich, which was then placed in an oven to cure at 60°C for 48 hrs. After polymerization, resin pieces containing gill arches were affixed with cyanoacrylate glue end-on to (i.e., thin edge of the sandwich facing) a blank resin block that had previously been trimmed with a Leica EM TRIM2 milling system. The sandwiches were glued at a 30–40° angle to the resin block to orient the gills, ensuring some lamellae were favorably aligned with the sectioning plane. This reduced the sectioning time required to capture entire lamellae.

#### Array tomography

After trimming the block face down, 60 nm serial sections were collected using an ATUMtome (RMC Boeckler) on glow-discharged Kapton tape (RMC Boeckler). Tape strips were cut and mounted onto 12 mm double-sided carbon tape with an aluminum base (EMS). The remaining sticky sides were then attached to 4-inch type-p silicon wafers (EMS) and grounded with conductive graphene carbon paint (EMS). Overview wafer images were collected using a Canon EOS 1300D camera. Wafers were placed in a Zeiss SEM (GeminiSEM 450; Carl Zeiss) and imaged using ATLAS5 Array Tomography software (Fibics). Six wafers were imaged using a 4-inch stage-decel holder and four-quadrant backscatter detector; the beam was operated at 4 kV EHT with 2 kV beam deceleration and 800 pA probe current. Low-resolution “street” images were collected at 3000 nm pixel sampling, medium-resolution section sets were collected at 150 nm pixel sampling, and high-resolution sites were collected at 15 nm pixel size. After image acquisition, the images were contrast inverted, cropped and aligned using the ATLAS 5 software. The image stack was exported and then processed using python-based scripts to produce a final 60 x 60 x 60 nm isotropic volume that was binned, aligned, and contrast/brightness adjusted.

#### FIB-SEM

The remaining AT specimen block was mounted on an SEM stub and sputter-coated with a thin layer of carbon (approximately 10nm, Leica ACE600). The specimen was transferred into the FIB-SEM instrument (Zeiss crossbeam 550) where Fields of View (FOVs) for vEM data collection were defined by acquiring an overview SEM image. Once target pillar cells were identified, an Atlas 3D (Fibics Inc., Ottawa) FIB sample preparation workflow was initiated. A 1 μm-thick protective platinum pad was deposited over the sample using an FIB current of 1.5 nA, after which tracking and autofocus lines were milled into the platinum surface and subsequently covered by an FIB-mediated deposition of 1 μm carbon at 1.5 nA. A coarse trench was milled using a 30 nA FIB beam and fine polished using a 3 nA beam. Milling and imaging parameters during the “continuous mill-and-image” acquisition run were set at 30 kV accelerating voltage, 1.5 nA current for the FIB, and 1.5 kV accelerating voltage, 1.1 nA beam current for the SEM. A pixel resolution of 5nm with a 15nm slice thickness was set, with a total dwell time of 2.6 us/pixel, and the EsB detector grid voltage was set at 820V. The imaging frame time was approximately 38 seconds, and the FIB advance rate was approximately 22.7nm/min. The raw image stacks were then registered, inverted, and binned using in-house scripts.

### Image Processing

Confocal images were processed using NIS Elements, with a subset of images being deconvolved using NIS Batch deconvolution. Maximum intensity projections of confocal stacks are shown for fluorescent confocal images. When required, time-lapse movies were aligned in XY using NIS Elements. Non-linear adjustments (gamma) were applied to some images to enhance the visualization of high-dynamic-range data. Time-lapse movies were generated in NIS Elements and exported to Adobe Premiere Pro CC 2024. Labels and arrows were added using Adobe Premiere Pro CC 2024 and Adobe Photoshop CC 2024. Schematics were created with Adobe Photoshop CC 2024, Microsoft PowerPoint, and BioRender.

FIB-SEM image stacks were processed using 3D Slicer software (v5.6.2). Multiple zAerocytes were identified in the sample by their morphology, and one was selected based on its placement within the sample and its standard, representative morphology. The “draw” feature was used to select the interior of a cell in an image and create a single slice of a 3D model of that cell. The process of selecting the interior of the cell and adding it to the model was repeated every 3-5 slices along the Z axis until the uppermost and lowermost images where the zAerocyte was visible. Then, the “fill between slices” feature was used to interpolate the rest of the structure, and the “smoothing” feature (Method: Gaussian, Standard Deviation: 35nm) was applied to reduce the impact of human variability on the surface shape of the cell. The same process was used to segment the interacting epithelial cells. The generated models were exported as TIF stacks and STL files for further analysis in Dragonfly 3D World software (v2024.1).

### Immunofluorescence

Dissected adult gills were fixed in 4% paraformaldehyde (PFA) prepared in 1x PBS at room temperature for 2 hours with gentle nutation. Following fixation, samples were washed in 1x PBS for 30 minutes, three times. Gill tissues were then incubated in Inoue blocking solution (5% normal goat serum; NGS, 2% BSA, 1.25% Triton X-100, in 1x PBS) (Inoue & Wittbrodt, 2011) for 1 hour at room temperature on a nutator. Incubation with primary antibodies was performed in I-buffer (1% normal goat serum, 2% BSA, 1.25% Triton X-100, in 1x PBS) for 2 days at 4°C on a nutator. Gill tissues were washed three times for 30 minutes each with 1x PBS at room temperature. Secondary antibodies, diluted in I-buffer, were applied for 1 day at 4°C on a nutator. After secondary antibody incubation, gill tissues were washed three times with 1x PBS for 30 minutes and kept at 4°C in the dark until imaging. Anti-serotonin antibody (Sigma-Aldrich; Cat# S5545-25UL; 1:500 dilution); Anti-neuronal cell surface marker (zn-12) (DSHB; Cat# supernatant 1.0 ml; 1:500 dilution); Phalloidin Far-Red (ThermoFisher; Cat# A22287); Anti-myosin light chain (phospho S20) (Abcam; Cat# ab2480; 1:500 dilution).

### Dye injections

Intravenous injections of DRAQ5^TM^ (ThermoFisher; Cat# 62254) and/or BODIPY Cell Tracker Green BODIPY (ThermoFisher; Cat# C2102) and/or Hoechst (ThermoFisher; Cat# H1399) were done using Drummond Nanoject III microinjector (Item# 3-000-207) with pulled glass capillary needles (Drummond item # 3-00-203-G/X) inserted under the scales in the ventral caudal region of the fish, into the axial vasculature. A volume of 100-150 nl was injected at a concentration of 1.7 mM for DRAQ5, 1.7ng/nl for BODIPY, and 1.6 mM for Hoechst.

### Preparation of zebrafish single-cell suspension for scRNA-seq

Adult zebrafish were euthanized by submersion in an ice bath. A pair of Dumont L5 forceps and surgical scissors were used to remove the gill basket. The gill arches were then further separated and dipped in 1x PBS to remove debris. The dissected gill arches were placed in Liberase (Sigma-Aldrich 05401119001). They were slowly pipetted up and down at room temperature for 45 minutes with a 1000 µL paipette set to 500 µL to ensure gentle dissociation. Cell dissociation was halted by adding an equal volume of STOP solution (5% FBS, 2% BSA in DMEM) (Corning 35-010-CV/Millipore Sigma A9418-5G/Gibco A1896701) and gently inverting the tube. The cell suspensions were filtered through a 40-micron filter and centrifuged at 500 × g for 5 minutes at room temperature. The supernatant was removed, and the cells were resuspended in a 1X PBS solution containing 2% BSA (Gibco 10010023/Millipore Sigma A9418-5G). The cells were counted on a LUNA-FL (Logos Biosystems) and diluted to an optimal concentration of 700 cells/uL using the same 1X PBS 2% BSA solution. Approximately 10,000 cells were loaded onto the 10x Genomics Chromium controller.

### Sequence alignment and quality control

Alignment of sequencing reads and processing into a digital gene expression matrix was performed using Cell Ranger version 7.0.0. The data were aligned against GRCz11 release 99 (January 2020) using the Lawson lab Zebrafish Transcriptome Annotation version 4.3.2 (Lawson et al., 2020). Cells were processed and analyzed using the latest Seurat (v4) (Satija et al., 2015) and R Studio. Cells with abnormally high (>2500) or low (<200) numbers of detected features, or with abnormally high mitochondrial content (>5%), were removed. The remaining cells were normalized to 10,000 transcripts per cell and scaled using the ScaleData function of Seurat with default settings. **Figure 2B** shows the cell and read count metrics.

**Figure 2.**
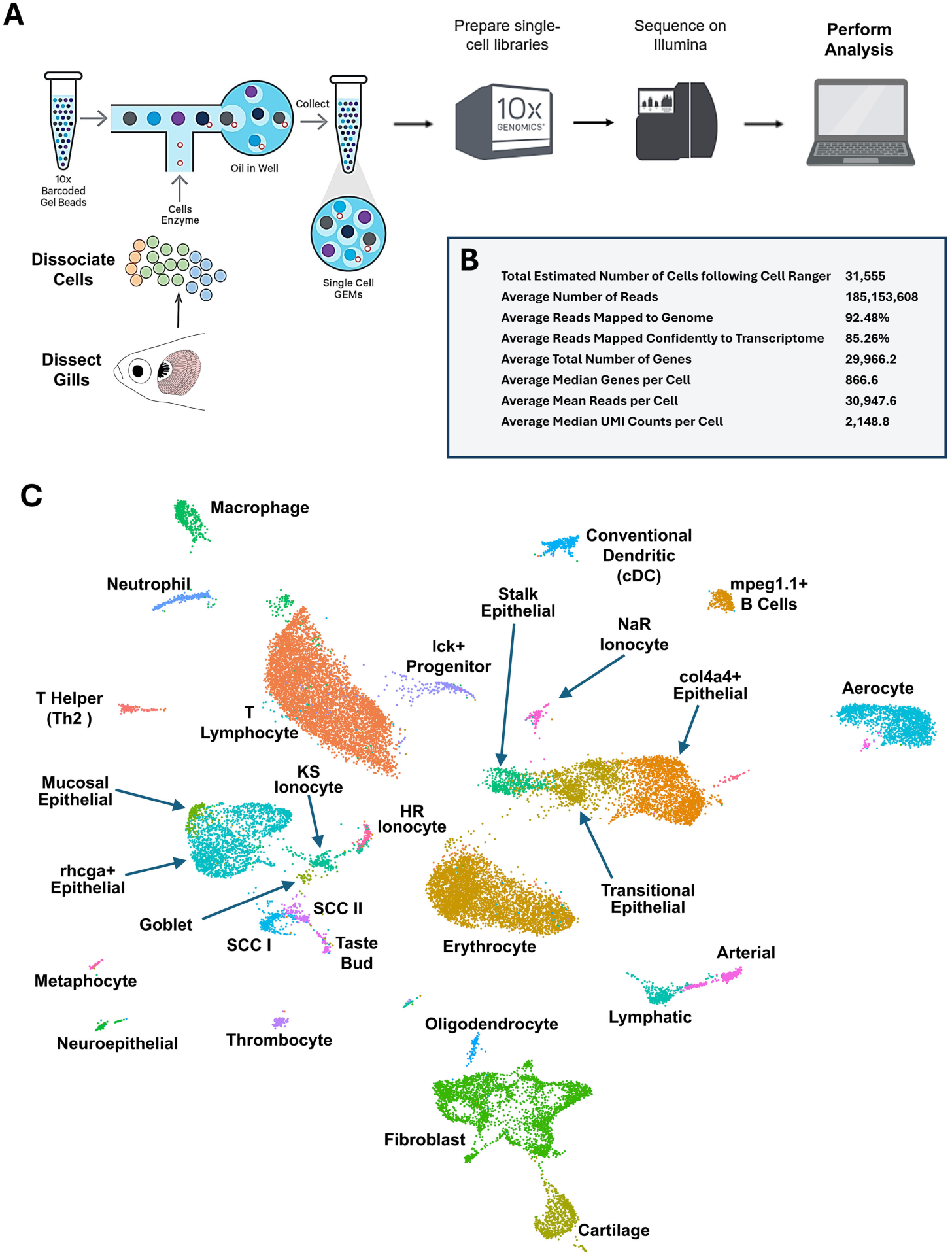
Single-cell RNA sequencing (scRNAseq) of the zebrafish gill. **A,** Schematic diagram showing the workflow for zebrafish gill scRNAseq. **B,** Metrics for the zebrafish gill scRNAseq. **C,** Uniform Manifold Approximation and Projection (UMAP) plot of data from the zebrafish gill scRNAseq, with 29 clusters annotated by cell identity.

### Dimensionality reduction, clustering, and visualization

The 2000 most variable features were identified for our sample using Seurat’s FindVariableFeatures function with default parameters. Principal component analysis (PCA) was conducted with Seurat::RunPCA, utilizing the most variable features determined earlier. Seurat::JackStrawPlot was used to identify the number of significant principal components for downstream analyses. Unbiased clustering was performed using Seurat::FindNeighbors and Seurat::FindClusters with default parameters. A Uniform Manifold Approximation and Projection (UMAP) was calculated using the RunUMAP function and visualized with DimPlot.

### Differential expression analysis

To identify differentially expressed genes across cell types, we used a negative binomial model implemented in the Seurat FindAllMarkers function, with a log fold change cutoff of 0.25. Genes were deemed differentially expressed if the adjusted P value was below 0.01.

### Defining cell types

Each of the 29 clusters was manually annotated based on an extensive survey of well-known tissue- and cell-type-specific markers. These markers were identified using various databases (The Zebrafish Information Network and Daniocell) and through an extensive literature search (Bradford et al., 2022; Sur et al., 2023). Our search and identification were guided by preliminary confocal imaging and electron microscopy of the zebrafish gill. For each cell type, at least three marker genes were identified.

### Hybrid cell doublet analysis

To simulate artificial epithelial-aerocyte doublets, we mixed gene expression signatures of the col4a4+ epithelial cluster (c3) and the aerocyte cluster (c39). For each of the two clusters, we then randomized the pool of c3 and c39 cells to create 5000 random cell pairs that we used for creating artificial doublets. We also randomly picked 5000 cells from the rest of the dataset (except c3, c39 and the hybrid cell cluster c40) to serve as a computational negative control. To create a mixed gene expression signature characteristic of doublets encountered in scRNAseq, for each cell pair between c3 and c39, the expression data was unlogged, the mean of the epithelial (c3) and aerocyte (c39) clusters’ expression was taken, and log expression was recalculated. We limited the genes used to calculate average expression to the 2000 highly variable genes calculated on the whole gill scRNAseq dataset. To understand how similar c40 is to the epithelial (c3) or aerocyte (c39) populations, we calculated Euclidean distances in variable gene expression space for cells within c40 (i.e., hybrid cells to hybrid cells), c40 to c3 (i.e., hybrid cells to col4a4+ epithelial cells), c40 to c39 (i.e., hybrid cells to aerocytes), between the hybrid cell population itself (c40) and the simulated doublets between c3 and c39 (i.e., hybrid cells to epithelial-aerocyte simulated doublets), and between hybrid cells and non-epithelial and non-aerocyte cells using the “dist” function in R. Cells in the c40 had shorter distances to the artificially simulated doublets that its distances to the epithelial (c3) and aerocyte (c39) clusters.

### Hybridization Chain Reaction (HCR)

HCR was performed to visualize and confirm the identification of cell types associated with the gill. HCR probe sets were designed by Molecular Instruments (Molecular Instruments, Los Angeles, CA) or custom-designed using the method described in (Kuehn et al., 2022) and ordered through IDT. Gills were dissected and fixed in 4% PFA (Electron Microscopy Sciences, Cat # 15710) prepared in 1X PBS for 2 hours at room temperature. They were then washed in 1X PBS (Gibco, 10010023) for 30 minutes, three times. Fixed gills were pre-hybridized with pre-heated (37 °C) HCR probe hybridization buffer (Molecular Instruments) for 30 minutes at 37 °C while rotating. Probe hybridization was performed with 2 μL of each 1 μM probe diluted in 500 μL of probe hybridization buffer at 37 °C, with rotation for 12 hours. The probe solution was removed by washing with preheated HCR probe wash buffer (Molecular Instruments) four times for 15 minutes each at 37 °C, followed by two 5-minute washes with 5X SSCT (12.5 ml 20X SSC in 50 µL Tween 20)(KD Medical Cat # RGF-3240/Millipore Sigma 9005-64-5) at room temperature before the pre-amplification stage. Gills were pre-amplified with HCR probe amplification buffer (Molecular Instruments) for 30 minutes at room temperature. Hairpins (H1 and H2) (10 μL of 3 μM stock) were pre-heated separately at 95 °C for 90 seconds and then cooled to room temperature for 30 minutes. Hairpins (H1 and H2) were then mixed with 500 μL of fresh HCR probe amplification buffer. The pre-amplification buffer was removed from the gills before adding the newly mixed hairpin solution, followed by a 12-hour incubation period in the dark at room temperature. Excess hairpins were then removed, and the samples were washed five times with 5X SSCT at room temperature. Samples were then stored at 4 °C in the dark until imaging.

## RESULTS

### Anatomy of the zebrafish gill

Like mammalian lungs, fish gills are complex, hierarchically organized organs with an enormous surface area dedicated to gas exchange (**Fig. 1A**). In lungs, gas exchange occurs primarily in the alveoli, tiny air sacs located at the end of the bronchioles. In fish, the analogous primary gas-exchange structures are gill lamellae. Zebrafish gills are located posterior to the eye and are externally situated underneath the operculum (gill cover). Zebrafish have four gill arches on either side of the head (**Fig. 1B**), each of which has 25-30 sets of paired feather-like gill filaments emerging perpendicularly from the arch (**Fig. 1C-F**). The gill arches also contain tooth-like gill rakers that help trap food and particulate matter from passing through and damaging the delicate gill filaments (**Fig. 1A, E, F**). The gills are readily accessible for high-resolution optical imaging and experimental manipulation, especially in individuals with naturally occurring opercular hypoplasia (**Fig. 1D**). Higher magnification imaging of individual zebrafish gill filaments shows that each filament has numerous rows of paired lamellae protruding from either side of the centrally located stalk (**Fig. 1G, H**). Histological sections reveal that lamellae are very thin, elongated leaf-like structures only about 10-20 µm thick, attached to wider central stalks with a cartilaginous core (**Fig. 1I, Supp. Fig. 1**). In *en face* views, lamellae are visible as leaf-like structures containing large numbers of circulating erythrocytes (**Fig. 1J, K**), as described further below.

### Single-cell analysis of the zebrafish gill

To characterize the cellular composition of the gills, we utilized the 10X Genomics Chromium platform to conduct single-cell RNA sequencing (scRNAseq) on single-cell suspensions prepared from four whole gill baskets (two from males and two from females) removed from adult zebrafish (**Fig. 2A**). An estimated total of 31,555 cells were sampled, with an average of 185,153,608 sequence reads; 92.5% of the reads were mapped to the genome, and 85.3% were mapped to the zebrafish transcriptome. This translates to an average of 30,948 sequence reads per cell, 2,149 median UMI counts per cell, and 867 median genes per cell (**Fig. 2B**). Unsupervised clustering using Seurat (Satija et al., 2015) resulted in 40 separate clusters that we annotated as 29 identifiable cell populations (**Fig. 2C, Supp. Fig. 2**) based on their expression of characteristic cell-type-specific genes, overall gene expression profiles, comparison to other single-cell RNAseq data sets, notably Daniocell (Sur et al., 2023), and spatial localization of cluster-specific transcripts to morphologically identifiable cells in the gills, as discussed below.

### Conserved cell types in the zebrafish gill

We identified several cell clusters corresponding to previously described gill cell types that have presumptive analogs in the mammalian lung (**Fig. 3A**). This was validated through comparisons to published scRNAseq databases, enriched expression of genes characteristic of each cell type (**Fig. 3B**), and spatial localization using *in situ* hybridization chain reaction (HCR) or available transgenic zebrafish lines (**Fig. 3C-O**). Clusters 3, 6, 9, 13, and 16 correspond to gill epithelial cell populations (**Fig. 3B**). Cluster 3 (*col4a4*+ epithelial cells) and cluster 16 (*rhcga*+ epithelial cells) localize to lamellar epithelium, as shown by HCR labeling for *collagen, type IV, alpha 4 (col4a4*; **Fig. 3C-D, P**) and *Rh family, C glycoprotein a (rhcga*; **Fig. 3E-F, P**), respectively. *rhcga*+ epithelial cells are more enriched on the deoxygenated edge of the lamellar “leaf” (**Fig. 3F, P**) while *col4a4+* epithelial cells appear more evenly distributed over the entire gill lamella (**Fig. 3D, P**). In humans, the *rhcg* gene is expressed in lung bronchial epithelial cells (Han et al., 2009), while type IV collagen plays an important role in driving alveolar epithelial-endothelial association in the lung blood-gas barrier (BGB) (Loscertales et al., 2016). Cluster 13 (stalk epithelial cells) shows specific expression of *cldna* and *notch3*, and HCR for *cldna* reveals localization to the epithelium surrounding the core (stalk) of the gill filament (**Fig. 3E-F, P**). In human lungs, *notch3* is expressed in suprabasal cells and helps determine the fate of epithelial cells during development as well as during repair and regeneration following lung injury (Bodas et al., 2022; Deprez et al., 2020; Hewitt & Lloyd, 2021; Mori et al., 2015). Cluster 6 shares many genes with cluster 3 (*col4a4*+ epithelial cells) and cluster 13 (*cldna*+ stalk epithelial cells) and appears between these two clusters in UMAP projections (**Fig. 3A, B**), suggesting that it is a transitional epithelial cell population at the stalk/lamella interface. Cluster 9 represents mucosal epithelial cells that are found on both the dorsal and ventral sides of gill filament stalks (**Fig. 3A,B,C-D, P**). Mucosal epithelial cells express muc13a, an ortholog of murine MUC13, which is found in pneumocytes and bronchiolar epithelium in the lung (Williams et al., 2001).

**Figure 3.**
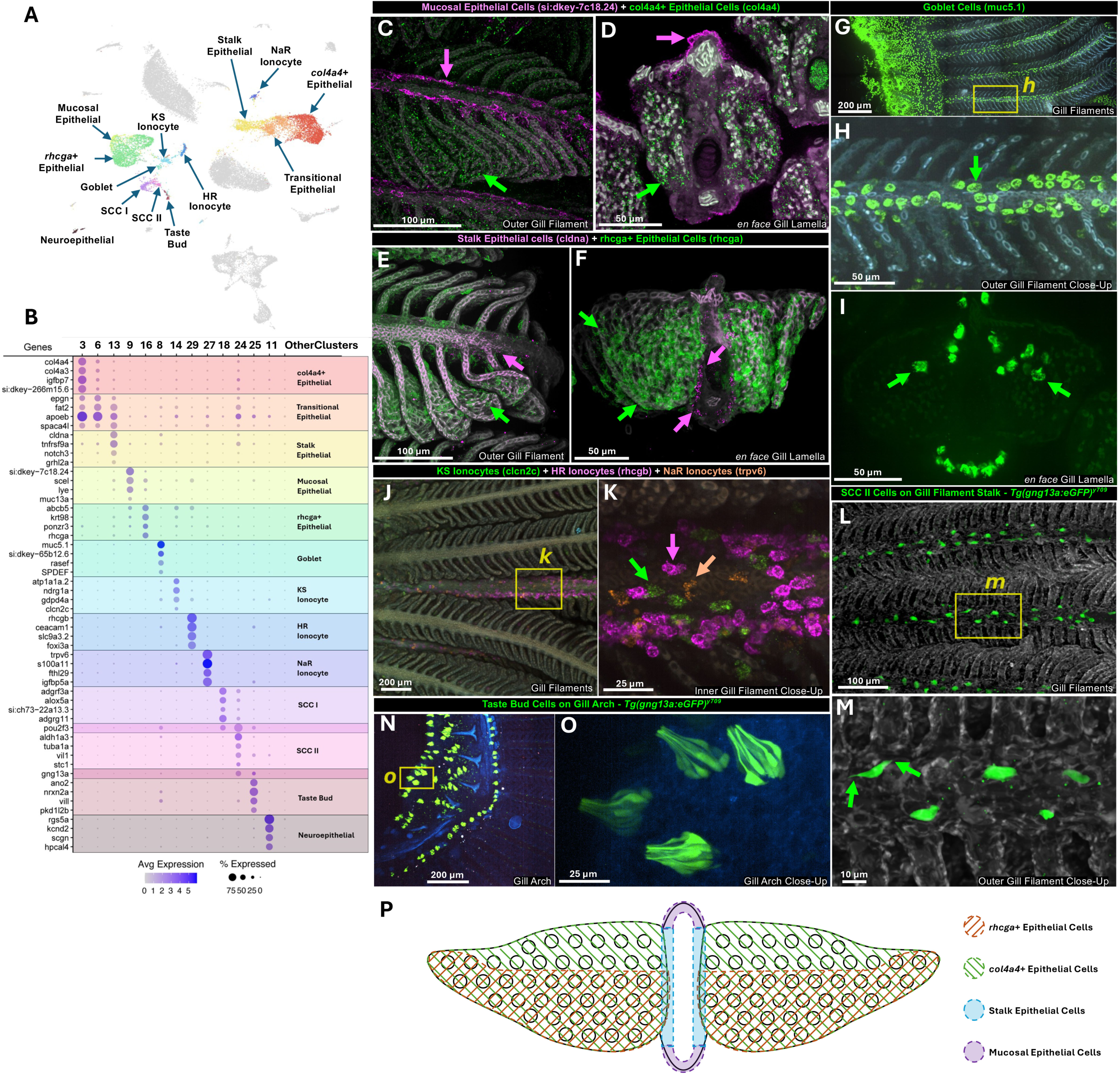
Epithelial cell types of the zebrafish gill. **A,** UMAP plot of zebrafish gill scRNAseq data highlighting 13 epithelial-related clusters. **B,** Dot plot showing the relative gene expression of genes used to identify and characterize the 13 clusters in panel A. **C,D,** Confocal micrographs of adult zebrafish gill subjected to hybridization chain reaction (HCR) for *si:dkey-7c18.24* (magenta) and *col4a4* (green), lateral view (C), and transverse section revealing *en face* gill lamellae (D). **E-F,** Confocal micrographs of adult zebrafish gill subjected to HCR for *cldna* (magenta) and *rhcga* (green), lateral view (E), and transverse section revealing *en face* gill lamellae (F). **G-I,** Confocal micrographs of adult zebrafish gill subjected to HCR for *muc5.1* (green), (G) overview showing gill arch and filaments, (H) higher magnification image of the boxed area in panel G, (I) transverse section revealing *en face* gill lamellae. **J,K,** Confocal micrographs of adult zebrafish gill subjected to HCR for *clcn2c* (green), *rhcgb* (magenta), and *trpv6* (orange), (J) overview showing multiple gill filaments, (K) higher magnification image of the boxed area in panel J. **L-O,** Confocal micrographs of gills dissected from an adult *Tg(gng13a:egfp)^y709^* transgenic zebrafish showing two different types of gng13a+ green fluorescent cells. (L) Overview showing multiple gill filaments with gng13a+ SCC II cells), (M) higher magnification image of the boxed area in panel L. (N) Overview of gill arch region showing gng13a+ taste bud cells, (O) higher magnification image of the boxed area in panel N. **P,** Schematic diagram summarizing epithelial cell localization in adult zebrafish gill lamellae based on specific marker gene expression; *en face* view. Scale bars = 100 µm (C,E), 50 µm (D,H,F,I), 200 µm (G,J,N), 25 µm (K,O), and 10 µm (M).

Cluster 8 corresponds to goblet cells (**Fig. 3A,B,G-I**). Goblet cells are found in both gills and lungs, where they protect epithelial surfaces by secreting mucus (Symmes et al., 2018). The thick, viscous mucus layer produced by these cells traps particles, pathogens, and debris, helping to prevent damage and infection. The previously characterized (Castranova et al., 2025) *mucin 5.1* (*muc5.1*) gene is highly enriched in the goblet cell cluster, as are a number of other genes (SPDEF) also found in mammalian lung goblet cells (**Fig. 3B**) (Chen et al., 2009). *In situ* HCR for *muc5.1* reveals goblet cells lining both the deoxygenated (inward-facing) and oxygenated side (outward-facing) surfaces of gill filament stalks (**Fig. 3G-I**). Transverse images of *en face* gill lamellae also show that a few goblet cells are present on the lamellar surface close to the oxygenated side of the filament stalks (arrows in **Fig. 3I**).

Clusters 14, 29, and 27 were readily identified as ionocytes, specifically KS, HR, and NaR ionocytes, respectively (**Fig. 3A,B,J,K, Supp. Fig. 3A,B**). KS ionocytes are distinguished by *clcn2c, atp1a1a.2,* and *ndrg1a* markers, and they perform Cl-uptake and HCO3-secretion. HR ionocytes specifically express *rhcgb, ceacam1, slc9a3.2,* and *foxi3a* and are required for Na+ uptake, H+ secretion, and NH4+ excretion. NaR ionocytes are distinguished by *trpv6* and *s100a11* markers, and they play an important role in Ca2+ uptake (Guh et al., 2015). Combined HCR for KS, HR, and NaR ionocytes using markers *clcn2c* (green), *rhcgb* (magenta), and *trpv6* (orange) reveals that all three types of ionocytes are found on the deoxygenated (inward-facing) side of the gill filament stalks (**Fig. 3J-K, Supp. Fig. 3A,B**). Ionocytes play an important role in osmoregulation in the fish skin and fish kidney by maintaining ionic balance, and also regulate acid-base balance and ammonia excretion (Dymowska et al., 2012; Hwang et al., 2011). In mammalian lungs ionocytes facilitate absorption of water and salt from the airway surface and help drain the liquid lining airways (Lei et al., 2023; Montoro et al., 2018; Plasschaert et al., 2018). One recent study reported three distinct populations of pulmonary ionocytes in the lungs of ferrets (Yuan et al., 2023), including type A pulmonary ionocytes expressing NKCC1 (slc12a2) and the voltage-gated chloride channel (clcnka), type B pulmonary ionocytes expressing the proton pump genes ATP6AP2 and ATP6V1F, and type C pulmonary ionocytes with enriched expression of aquaporin genes (AQP3, 4, and 5). Preliminary comparison suggests type A most closely resembles the zebrafish *clcn2c+* KS ionocytes while type B resembles zebrafish *atp6ap2+* and *atp6v1f+* HR ionocytes.

Clusters 18 and 24 are *pou2f3*-expressing gill solitary chemosensory cells (SCCs), known as Tuft cells or brush cells in lungs (**Fig. 3A,B)**. In lungs, these cells reside in the respiratory airways where they play a role in chemosensation, antigen detection, and responses to other stimuli (Barr et al., 2022; Billipp et al., 2021; Li et al., 2022). HCRs for *adgrf3a* and *aldh1a3* label SCC I (cluster 18) and SCC II (cluster 24) cells, respectively. SCC I cells localize sparsely all around gill filament stalks while SCC II cells are found mainly on the deoxygenated (inward-facing) side of the stalk (**Supp. Fig. 3C,D**). SCC II (cluster 24) and taste bud chemosensory cells (cluster 25; **Fig. 3A,B**) both express *guanine nucleotide binding protein (G protein), gamma 13a* (*gng13a*), and are readily visualized using a *Tg(gng13a:egfp)^y709^* transgenic line (Castranova et al., 2025). Transgene-positive SCC II cells on the gill filament shaft (**Fig. 3L,M**) resemble the “taste-like” solitary chemosensory type 2 (SCC2) cells reported in the recently described zebrafish Axillary Lymphoid Organ (Castranova et al., 2025), including the presence of paired apical protrusions on opposite ends of the cell (**Fig. 3L-M**; arrows in M). Transgene-positive taste chemosensory cells with “taste bud-like” morphology are found on the gill arches and gill arch rakers (**Fig. 3N-O**).

Cluster 15 includes neuroepithelial cells (NECs) of the gill (**Fig. 3A-B**). Gill NECs and mammalian pulmonary neuroendocrine cells (PNECs) are strikingly similar cells that detect changes in oxygen levels and initiate physiological responses. Evidence for the role of gill NECs in oxygen chemoreception has come from whole-cell patch-clamp studies in the zebrafish (Jonz et al., 2004). Gill NECs and lung PNECs also share signaling molecules, including 5-HT (serotonin) (Candeli & Dayton, 2024; Pan et al., 2022) and even an endodermal origin (Hockman et al., 2017). Immunofluorescence using anti-serotonin antibody localizes gill NECs to the gill filament stalk (**Supp. Fig. 3E,F**), as previously reported (Jonz et al., 2004). Together, these findings show zebrafish gills possess a number of key cell types that resemble those found in mammalian lungs, both in terms of cell origin, gene expression, and function.

### Vascular architecture of the zebrafish gill

Like mammalian lungs, fish gills possess a highly complex vascular network with a vast surface area facilitating gas and ion exchange. Deoxygenated blood pumped from the fish heart is oxygenated in the gills before being directed to the rest of the body of the fish (**Fig. 4A**). The four gill arches each contain a single large afferent branchial artery carrying deoxygenated blood from the heart into the gill arch, and a single efferent branchial artery that carries oxygenated blood from the gills to the dorsal aorta. Both sets of branchial arteries branch into filamental arteries at the base of each of the gill filaments (**Fig. 1A, 4B**). Blood flows from the afferent branchial artery into afferent filamental arteries running proximal-to-distal through the filament stalks, from where it is directed into the numerous flattened lamellar leaflets projecting laterally from the gill filament stalks (**Fig. 4B,C**). After circulating through lamellar leaflets oxygenated blood exits the other side of the base of the lamella and enters the efferent filamental artery running distal-to proximal through the other side of the filament stalks before draining into the efferent branchial artery for circulation to the rest of the body (**Fig. 4C**). The complex zebrafish gill vasculature can be visualized using a *kdrl* transgenic line (**Fig. 4D-H, Supp. Movies 1,2**) and other vascular reporter lines (see below). Highly vascularized gill filaments emerge at right angles from the gill arches (**Fig. 4D-E**), with numerous *kdrl:mCherry-*positive lamellae extending from each gill filament shaft at regular intervals (**Fig. 4F-H**). Robust circulation through the filament stalks and lamellae can be easily visualized using either transmitted light or confocal imaging of animals with fluorescently “tagged” blood vessels and blood cells (**Fig. 1K, Supp. Movies 1,2**). Electron microscopic array tomography of an isolated adult zebrafish gill filament and lamella reveals the ultrastructural morphology of its vascular spaces in detail (**Fig. 4I-K, Supp. Fig. 4, Supp. Movie 3**). Afferent and efferent filamental arteries are both readily visible in transverse sections through the gill filaments, where they appear as erythrocyte-filled tubes on either side of the lamellar stalk (**Fig. 4I-K, Supp. Fig. 4A,B,D**). In other sections in the array tomography volume the filamental arteries can be seen branching into the rims of the lamellae (**Supp. Fig. 4A,C,D**), and erythrocytes can be seen threading between cells with irregularly shaped nuclei within the lamellar leaflets themselves (**Supp. Fig. 4E,F**), as is also seen in live images of erythrocytes circulating through lamellae (**Supp. Movie 2**). Segmentation and 3-D reconstruction of the luminal spaces, erythrocytes, and epithelial layers from the array tomography dataset further highlights the path of flow between filamental arteries through the lamellar leaflet. (**Supp. Movie 3**). Although confocal imaging of mature lamellae using *Tg(kdrl:mcherry)^y206^* or *Tg(fli1a:eGFP)^y1^* transgenic lines reveals strong transgene expression in endothelial cells within the filament shaft or lining the lamellar rim, these transgenes are only very weakly expressed in cells in the center of the lamellar leaflets (**Fig. 4H, Supp. Fig. 5A-F, Supp. Movie 2**), although stronger *Tg(fli1a:eGFP)^y1^* expression is visible in cells in the center of immature lamellae located at the distal tips of gill filaments (**Supp. Fig. 5G,H**).

**Figure 4.**
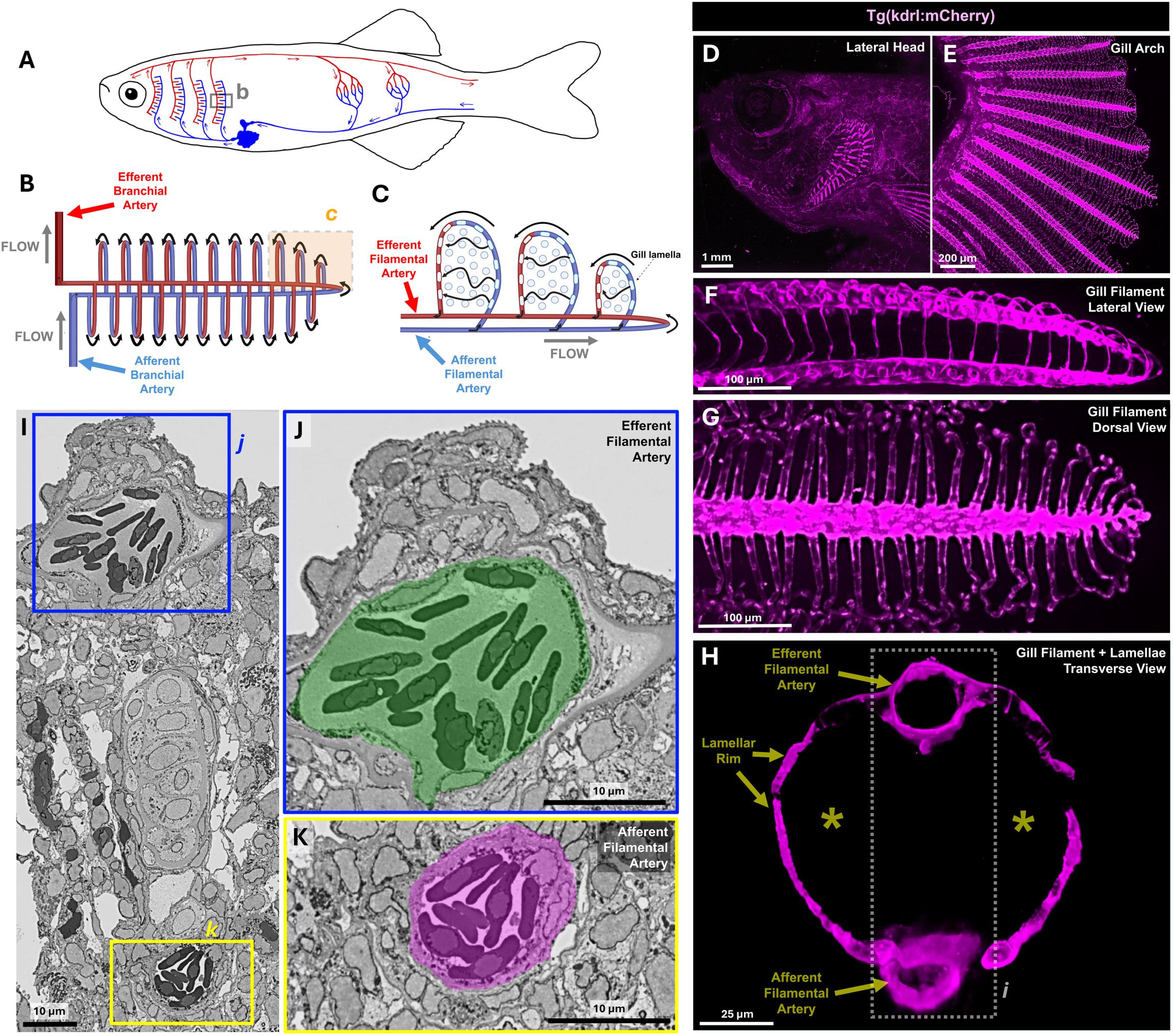
Vascular anatomy of the zebrafish gill. **A-C,** Schematic diagrams illustrating (A) simplified vascular anatomy of blood flow through gas-exchange vasculature and body of an adult fish, (B) higher magnification view of blood flow through an individual gill filament (boxed area in panel A), and (C) even higher magnification view of circulation through individual gill lamellae (boxed area in panel B). **D-H,** Confocal micrographs of adult *Tg(kdrl:mCherry)^y206^* transgenic zebrafish showing (D) lateral head view, (E) dissected gill arch, (F) lateral view of gill filament, (G) dorsal view of gill filament, and (H) *en face* view of gill lamellae. See also **Supp. Movies 1,2** for comparable live images from adult *Tg(kdrl:mCherry)^y206^* transgenic zebrafish. The boxed area in H corresponds to the approximate area shown in panel I. **I-K,** Individual array tomography electron micrograph transverse sections through a gill filament, revealing the filamental arteries. The boxed areas in (I) are magnified in panels (J) and (K), showing the oxygenated efferent filamental artery and deoxygenated afferent filamental artery. See also **Supp. Movie 3** for reconstructions from the entire array tomography volume. Scale bars = 1000 µm (D), 200 µm (E), 100 µm (F,G), 25 µm (H), and 10 µm (I-K).

**Figure 5.**
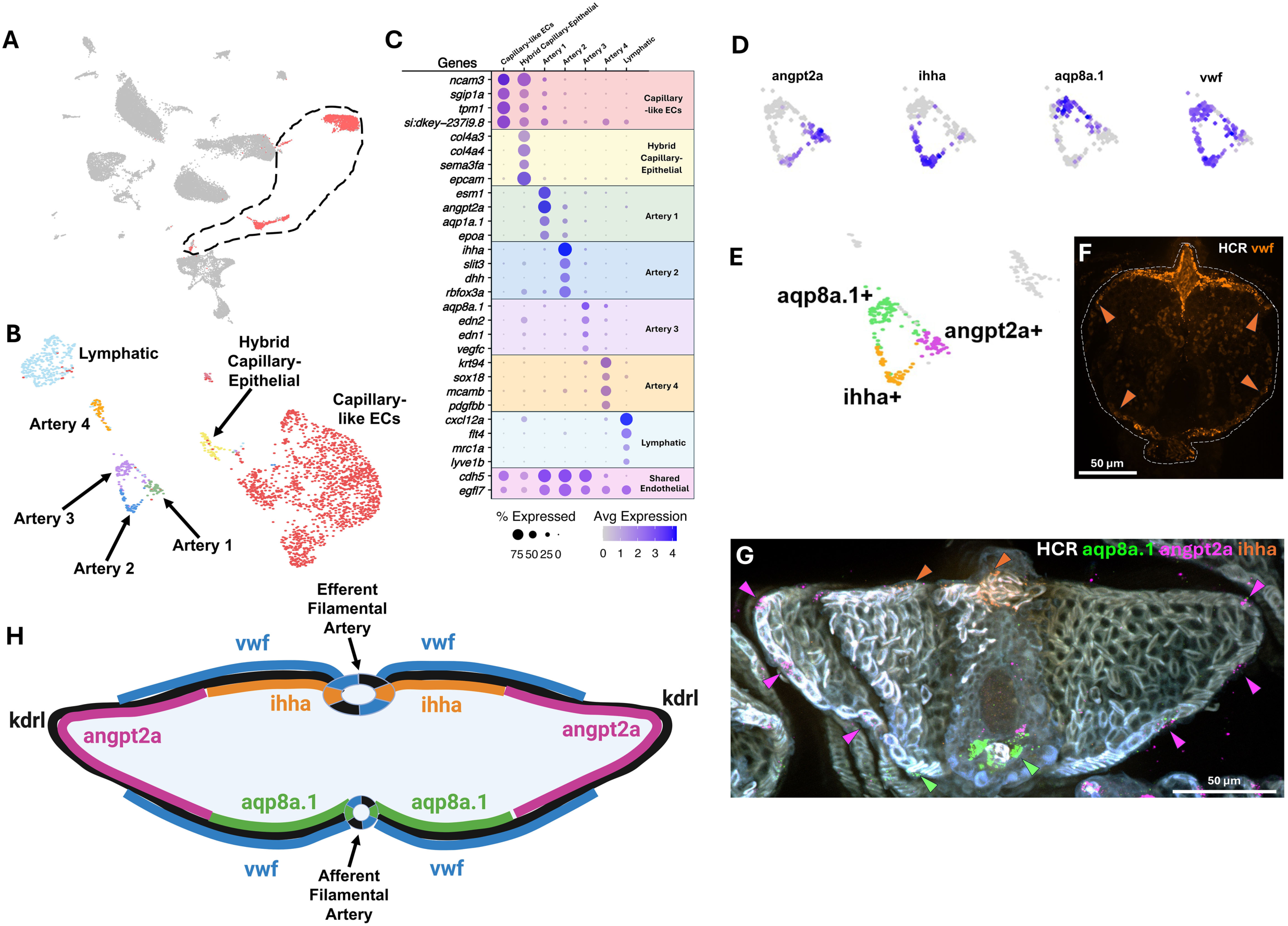
Endothelial cell types and arterial heterogeneity of the zebrafish gill. **A,** Uniform Manifold Approximation and Projection (UMAP) plot of data from the zebrafish gill scRNAseq, with endothelial cells selected for subclustering in panels B and C circled and highlighted in red. **B,** UMAP plot of subclustered endothelial cells highlighted in panel A. **C,** Dot plot showing the relative gene expression of genes used to identify and characterize the in panel B. **D,** Feature plot of gene expression in the Artery 1, 2, and 3 subclusters identified in panels B and C. **E,** Summary plot of expression of genes in panel D. **F,G,** Transverse *en face* view confocal micrographs of adult zebrafish gill filaments subjected to HCR for *vwf* (F) or *aqp8a.1*, *ihha*, and *angpt2a* (G). **H,** Schematic diagram summarizing the localized expression of different marker genes in the gill arterial endothelium. Scale bars = 50 µm (F,G).

### Vascular heterogeneity of the zebrafish gill

To further explore the identity of gill endothelial cell populations, we used *cdh5* and/or *egfl7* expression to select endothelial cells from our full adult gill scRNAseq data set (**Fig. 5A**) and then re-clustered only the endothelial cell subset to better identify the specific subpopulations present (**Fig. 5B**). Seven separate clusters were uncovered, including 4 distinct clusters of *kdrl+* arterial ECs, an *mrc1a+* lymphatic EC cluster, a large *ncam3*+ capillary-like EC cluster, and a cluster expressing markers of both *ncam3+* endothelial and epithelial identity (**Fig. 5B,C**). Artery clusters 1-3 can be distinguished by their enriched expression of *angpt2a*, *ihha*, and *aqp8a.1*, respectively (**Fig. 5D**), with the three clusters forming a unique ring-like arrangement in the UMAP projection (**Fig. 5E**). The *vwf* gene is expressed throughout most of this “ring” except for a small part of the angpt2a expression domain (**Fig. 5D,E**). *In situ* HCR using probes targeting each of these genes reveals that their arrangement in the UMAP projection (**Fig. 5D,E**) mirrors their distinct spatial localization in gill filamental arteries and lamellar leaflets (**Fig. 5F-H**). Specifically, *aqp8a.1* localizes to the deoxygenated afferent filamental arteries and adjacent portions of the lamellar rim, *ihha* localizes to the oxygenated efferent filamental arteries and adjacent portions of the lamellar rim, *angpt2a* localizes to the distal edges of the lamellar rim, while *vwf* marks all of the *kdrl+* vasculature except for a portion of the *angpt2a* expression domain at the distal tips of the lamellar leaflets (**Fig. 5F-H**).

A lymphatic endothelial cluster is also readily identifiable based on its expression of characteristic “lymphatic genes” such as *mrc1a*, *flt4*, and *lyve1b* (**Fig. 5C**, **Fig. 6A-C**). Imaging of the gills in *Tg(mrc1a:eGFP)^Y251^*, *Tg(kdrl:mcherry)^y206^* double transgenic animals reveals a complex, loosely organized network of mrc1a+, kdrl-lymphatic vessels running down the center of the gill filament stalk between the two mrc1a-, kdrl+ filamental arteries (**Fig. 6D-H, Supp. Fig. 6A,B, Supp. Movie 4**). Array tomography confirms the presence of endothelial-lined lymphatics surrounding the cartilaginous spine of the filament stalk, with an occasional white blood cell observed within the lumen (**Supp. Fig. 6C-H, Supp. Movie 5**). Confocal imaging with additional lymphatic reporter lines, including *Tg(flt4:YFP)^hu4881^* (**Fig. 6I**) and *Tg(5.2lyve1b:DsRed)^nz101^* (**Fig. 6J**), confirms the lymphatic identity of these central stalk vessels.

**Figure 6.**
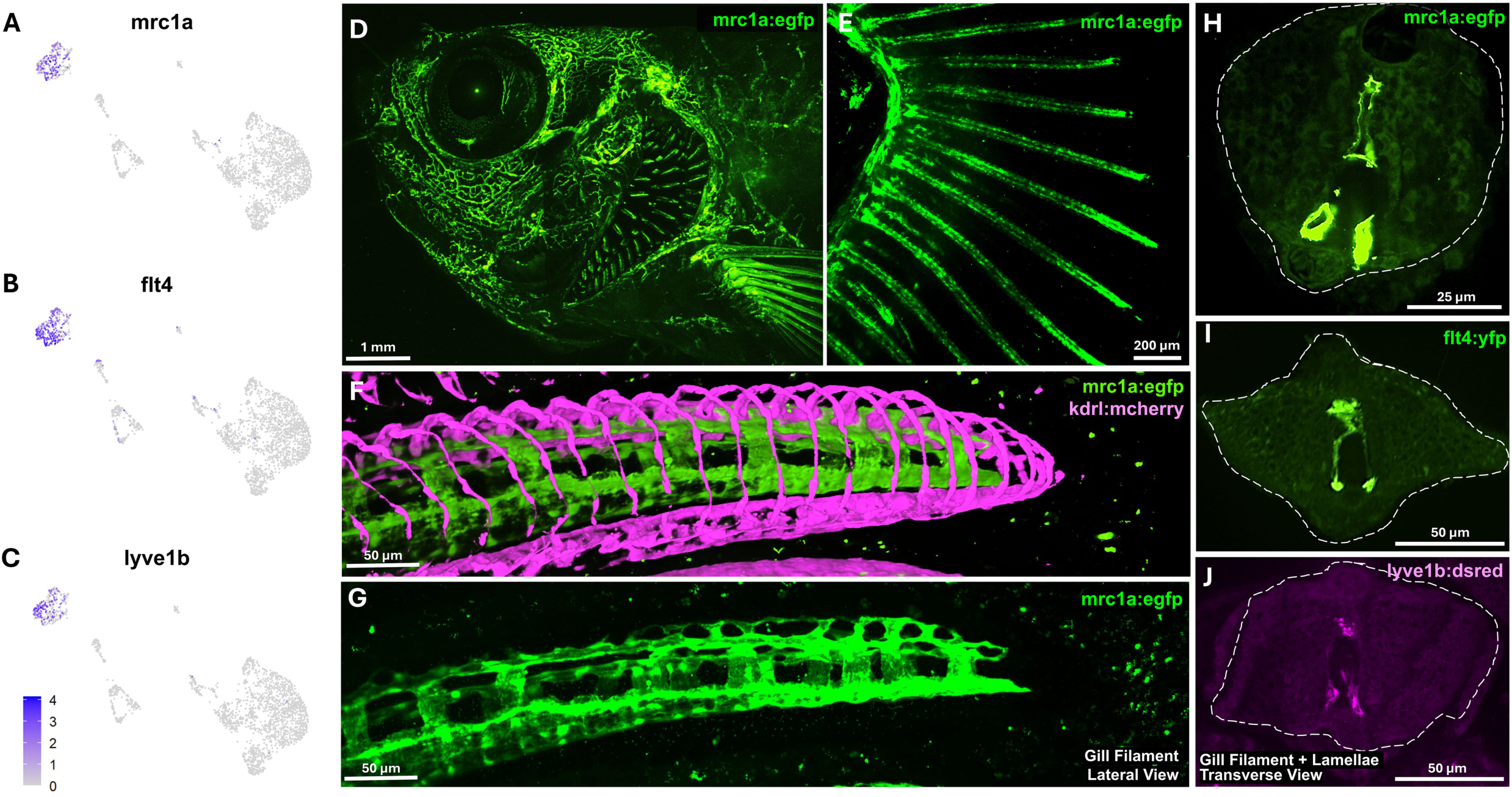
Lymphatic vasculature of the zebrafish gill. **A-C,** Feature plots of lymphatic gene markers on subclustered endothelial cells from Fig. 5B. **D-H,** Confocal micrographs of adult *Tg(kdrl:mcherry)^y206^* (arteries); *Tg(mrc1a:egfp)^y251^* (lymphatics) double transgenic zebrafish showing (D) a lateral view of the head (EGFP only), (E) a dissected gill arch (EGFP only), (F) a lateral view of the tip of an individual gill filament (EGFP and mCherry), (G) a lateral view of the same tip of an individual gill filament (EGFP only), and (H) transverse *en face* view of an individual gill filament (EGFP only). See also **Supp. Movie 4** for 3D reconstructions of the data from panels F and G. **I,J,** Transverse *en face* view confocal micrographs of adult zebrafish gill filaments from (I) *Tg(flt4:yfp)^hu4881^* or (J) *Tg(5.2lyve1b:dsred)^nz101^* transgenic zebrafish. Scale bars = 1 mm (D), 200 µm (E), 50 µm (F,G,I,J), 25 µm (H).

The largest gill endothelial scRNAseq cluster contains capillary-like *ncam3*+ endothelial cells (**Figs. 5B,C**, **Fig. 7A**). The single-cell transcriptional profile of these cells resembles that of mammalian Cap2 or “Aerocyte” endothelial cells, including enriched expression of zebrafish orthologs of many “diagnostic” genes highly expressed in murine Aerocyte (Cap2) cells (**Fig. 7B**), but not genes strongly expressed in lung “general capillary (gCap, Cap1) cells (Ellis et al., 2020; Gillich et al., 2020; Niethamer et al., 2020) (**Supp. Fig. 7A**). Whole mount *in situ* HCR of adult zebrafish gills shows that *ncam3*+ is expressed in the largely *Tg(kdrl:mcherry)^y206^* and *Tg(fli1a:eGFP)^y1^* negative cells in the center of the gill lamellae (**Fig. 7C-E, Supp. Fig. 7B-F, Supp. Movie 6**). These cells represent the major endothelial cell type of gill lamellae, populating all but the lamellar rim. Endothelial cells lining the outermost edge of the lamellar rim do not express *ncam3*, nor do the filamental artery or lymphatic endothelial cells found within the filament stalks, as described above. Based on their shared gene expression with mammalian Aerocytes and their comparable localization as the major endothelial cell population present in the primary gas-exchange structures of gills and lungs (lamellae and alveoli, respectively), we provisionally refer to these *ncam3*+ capillary cells as “zebrafish Aerocytes,” or “zAerocytes” for the remainder of this study (**Fig. 7E**).

**Figure 7.**
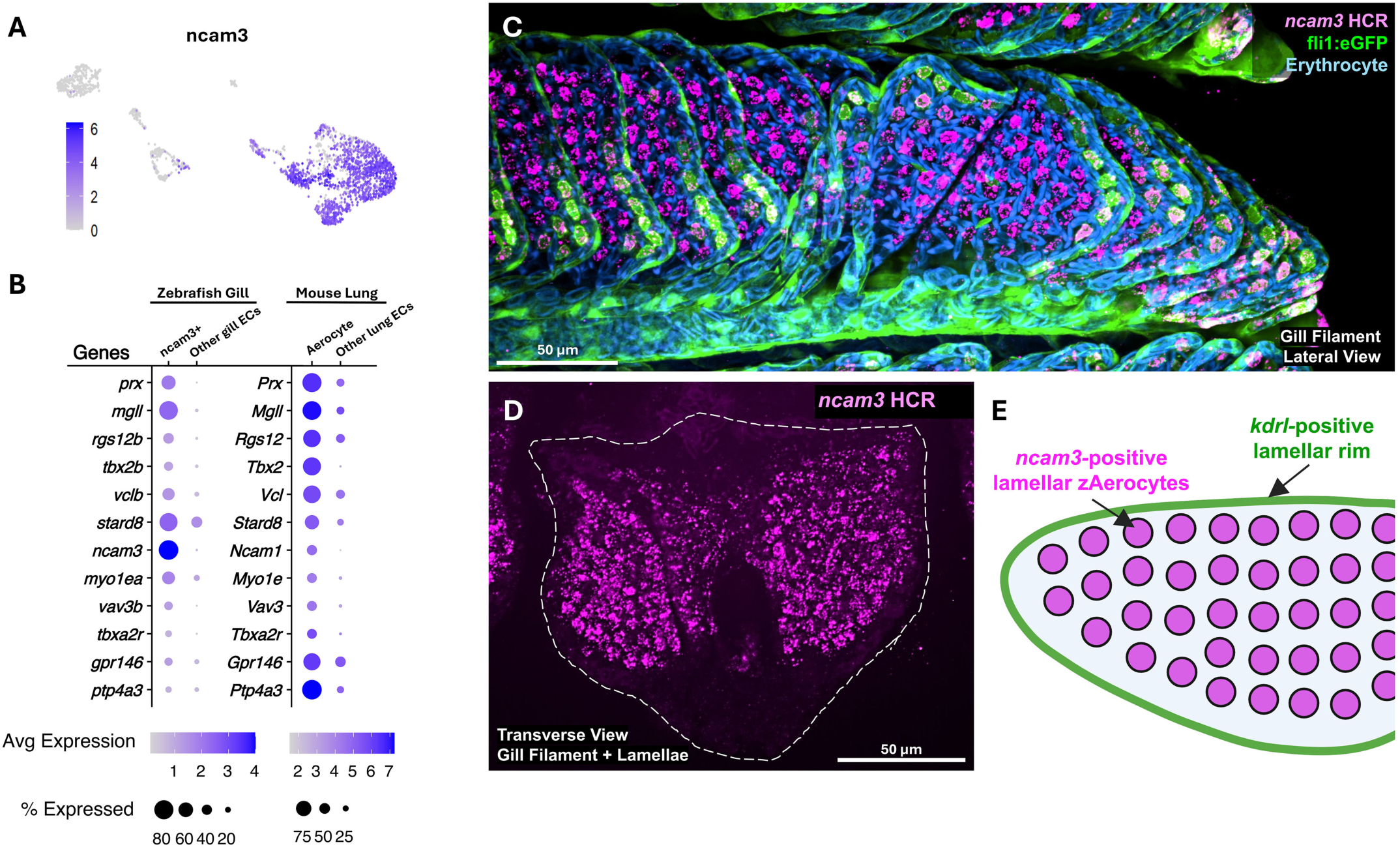
Capillary endothelial cells of zebrafish gill lamellae. **A,** Feature plot of *ncam3* expression on subclustered endothelial cells from Fig. 5B. **B,** Dot plots comparing gene expression in zebrafish gill *ncam3+* ECs or mouse lung Aerocytes with other endothelial cells identified in each of these two tissues, respectively. **C,** Lateral view confocal micrograph of the tip of an individual gill filament from an adult *Tg(fli1a:eGFP)^y1^* (green) animal, subjected to HCR for *ncam3* (magenta), with autofluorescent erythrocytes (blue). See also **Supp. Movie 6** for 3D visualization of the data from panel C. **D,** Transverse *en face* view confocal micrograph of an adult zebrafish gill filament subjected to HCR for *ncam3*. **E,** Schematic diagram of a gill lamella showing kdrl+, ncam 3-endothelial cells along the rim of the lamella, and kdrl-, *ncam3*+ zAerocytes in filling the central portion of the lamella. Scale bars = 50 µm (C,D).

### Morphology of zAerocytes

We used standard transmission electron microscopy (TEM), array tomography, focused ion beam scanning electron microscopy (FIB-SEM), and high-resolution confocal imaging to explore further the morphological characteristics of zAerocytes (**Fig. 8**). Transverse TEM sections though gill lamellae reveal erythrocyte-containing luminal spaces lined by a very thin endothelium, with the endothelium surrounded by a thin epithelial layer exposed to the outside environment (**Fig. 8A,B**). As in lung alveoli, the thin, closely juxtaposed endothelial and epithelial cell layers of the gill lamellae provide a short transcellular path with a large surface area, facilitating gas and ion exchange between the vascular lumen and the external environment (**Fig. 8B**). zAerocyte cell bodies, mostly filled by a large nucleus, are centrally located within the lamellar leaflet between luminal spaces (**Fig. 8A**). Complete ultrastructural reconstruction of an individual zAerocyte using FIB-SEM confirms its “wheel hub-like” shape (**Fig. 8C,D, Supp. File 1**). Red blood cells circulate around and past the ncam3+ zAerocyte cell bodies (**Fig. 8E-G, Supp. Movie 7**).

**Figure 8.**
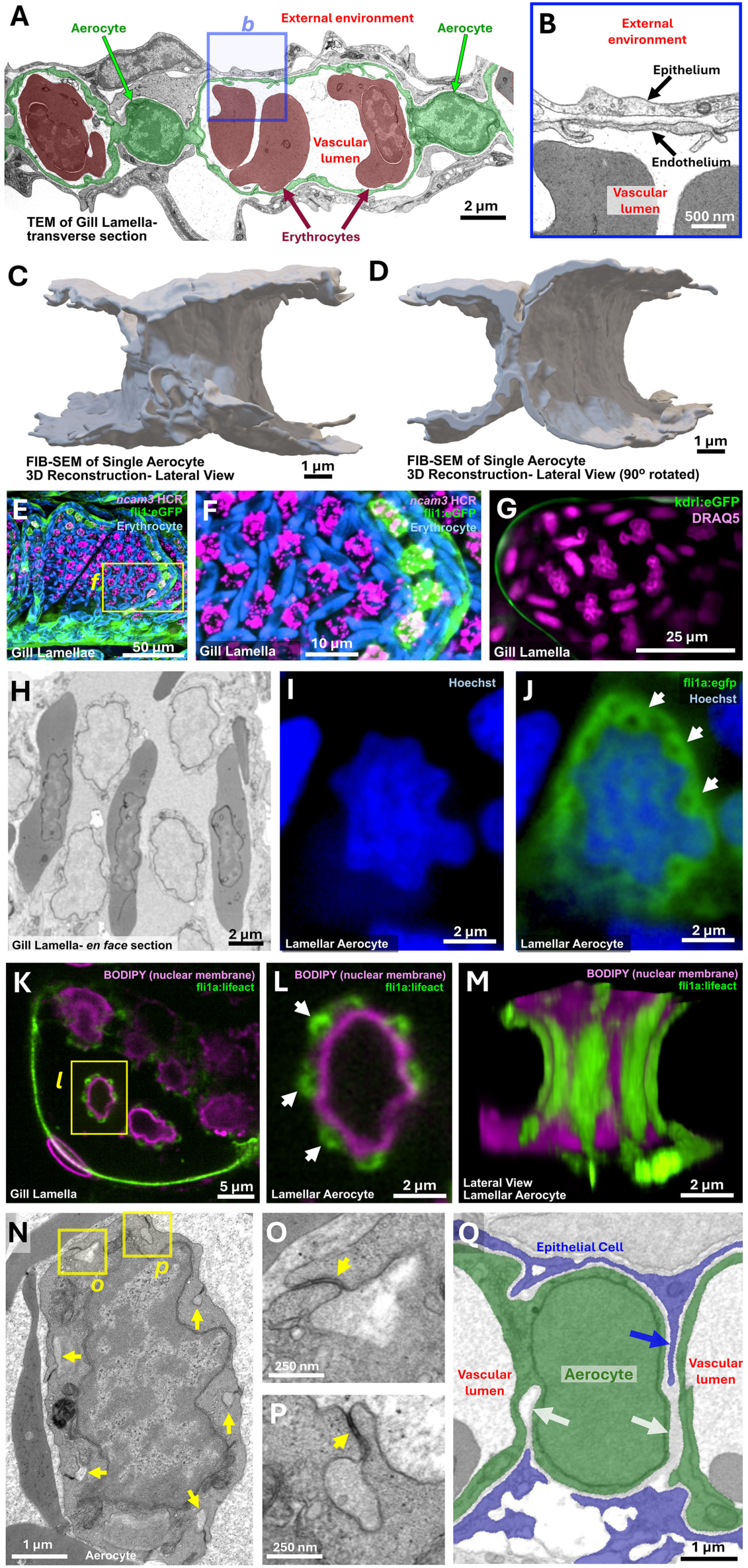
Morphology and ultrastructure of zebrafish Aerocytes. **A,B,** Standard TEM micrographs showing transverse sections through a gill lamella. (A) Two lamellar aerocytes (pseudocolored green) form enclose a vascular lumen containing erythrocytes (pseudocolored red). Box in panel A shows area magnified in panel B. (B) Higher magnification view of the thin gas-exchange epithelium and endothelium separating the external environment from the lamellar vascular lumen. **C,D,** Focused ion beam scanning electron microscopy (FIB-SEM) 3D segmentation and reconstruction of a single zebrafish Aerocyte showing a (C) lateral view, and (D) 90° rotated lateral view. **E,F,** Confocal micrographs of an adult *Tg(fli1a:eGFP)^y1^* transgenic zebrafish gill filament subjected to HCR for *ncam3* (magenta), with autofluorescent erythrocytes (blue). Box in panel E shows area magnified in panel F. Images are details from Fig. 7C. **G,** Confocal micrograph of an individual gill lamella in a living adult *Tg(kdrl:mcherry)^y206^* transgenic zebrafish gill lamellae after intravascular injection of nuclear dye DRAQ5. DRAQ5 labels both the stationary *ncam3*+ zAerocyte and circulating erythrocyte nuclei (teleost erythrocytes are nucleated). See also **Supp. Movie 7** for live images of the same lamella shown in panel G. **H,** Individual array tomography electron micrograph from an *en face* section through a gill lamella, showing erythrocytes threading through the lamellar lumen between aerocytes with large, irregularly shaped nuclei. **I,J,** Confocal micrographs of an individual aerocyte in an adult *Tg(fli1a:eGFP)^y1^* transgenic (green) zebrafish lamella stained with nuclear Hoechst dye (blue), showing Hoechst alone (I) or GFP + Hoechst together (J). White arrows in J point to “pillars.” **K-M,** Confocal micrograph of an individual lamella in an adult Tg(fli1a: LifeAct-eGFP) transgenic (actin; green) animals after intravascular injection with BODIPY546 (nuclear membranes; magenta). Images show (K) *en face* overview of the tip of the lamella, with a box noting the area magnified in panel L, (L) magnified image of a single aerocyte (single confocal plane from the center of the cell), and (K) 90-degree rotated side view of the aerocyte in panel L. White arrows in L point to actin-rich “pillars.” See also **Supp. Movie 8** for 3D visualization of the data from panels K-M. **N-P,** (N) Standard TEM micrograph from an *en face* section through a gill lamella, showing an individual aerocyte with prominent irregularly shaped nucleus and hollow “pillars” running through the adjacent cytoplasm (yellow arrows). Boxes in panel N show area magnified in panels O and P. (O,P) magnified images of individual aerocyte “pillars” sealed off from the lamellar lumen by autocellular junctions (yellow arrows). **Q,** Individual FIB-SEM section segmented to show epithelial cells (blue) and aerocytes (green) showing interaction between Aerocytes and lamellar epithelial cells. Aerocytes are penetrated both by “empty” extracellular spaces (white arrows) as well as by thin epithelial cell protrusions (blue arrow). Scale bars = 2 µm (A,H-J, L,M), 500 nm (B), 1 µm (C,D,N,Q), 50 µm (E), 10 µm (F), 25 µm (G), 5 µm (K), and 250 nm (O,P).

At higher magnification, zAerocyte nuclei are seen to be highly irregular in shape (**Fig. 8G-I, Supp. Movie 7, Supp. Fig. 4E,F**). These nuclei are surrounded by unusual, closely apposed actin-rich, hollow-cored “pillars” that run from top to bottom of the cells, often located in indentations or grooves in the sides of the nuclei (**Fig. 8I-M, Supp. Fig. 8**), accounting for previous reports that described these cells as “pillar cells” (Evans et al., 2005; Wilson & Laurent, 2002). The actin structures associated with the pillars are present intracellularly within the zAerocytes, as shown by their visualization using an endothelial-specific *Tg(fli1a:lifeact)^mu240^* transgenic line (**Fig. 8K-M, Supp. Fig. 8G-O, Supp. Movie 8**), but the central “holes” they surround (e.g., arrows in **Fig. 8J,L**) are matrix-filled extracellular spaces that generally run entirely through the zAerocyte from dorsal to ventral, as revealed using TEM and FIB-SEM (**Fig. 8N-Q, Supp. Fig. 9, Supp Movie 9**). As these extracellular spaces pass through the zAerocyte cell, these extracellular “pillar holes” are sealed off from the surrounding vascular lumens of the lamella by zAerocyte autocellular junctions (**Fig. 8O,P**).

zAerocytes are frequently deeply penetrated by thin cellular protrusions from adjacent, closely associated epithelial cells (**Fig. 8Q, Supp. Fig. 10**), although we have not observed individual epithelial cells completely bridging zAerocytes or making direct contact with epithelial cells from the other face of the lamella in our FIB-SEM data set. Interestingly, a small “hybrid zAerocyte-epithelial” cluster is present in our scRNAseq data with both aerocyte and col4a4+ epithelial gene expression signatures (**Fig. 5B,C, Supp. Fig. 11A-C**). Given the extremely close association between aerocytes and lamellar epithelial cells we hypothesized that this cluster might represent epithelial contamination of aerocytes or aerocyte/epithelial “doublets.” To validate our hypothesis, we computationally simulated doublets between col4a4+ epithelial cells (cluster 3) and zAerocytes (cluster 39) and estimated their transcriptomic similarity to the “hybrid” cells (cluster 40). We observed that the endogenous hybrid cluster is more similar to computationally simulated col4a4+ epithelial-zAerocyte “doublets” than it is to either the col4a4+ epithelial or zAerocyte cluster alone or to any other cluster in the scRNAseq data, and only slightly less similar than the hybrid cluster is to itself (**Supp. Fig. 11D**). Genes that are highly expressed in the simulated doublets were also highly expressed in the endogenous hybrid cluster. Furthermore, the endogenous hybrid cluster does not exhibit a unique gene expression signature; all markers strongly expressed in this cluster are shared with either col4a4+ epithelial or zAerocyte cells, suggesting that this cluster does not represent a unique cell subpopulation.

## DISCUSSION

In this study, we have used a variety of anatomical and molecular methods to examine the structure and cellular composition of the zebrafish gills and their vasculature. Gills are complex gas-exchange organs that share many features with mammalian lungs, including an intricate, highly ramified vascular network. Unlike mammalian lungs, however, the gills are externally located and easily accessible for high-resolution optical imaging and experimental manipulation, making them an excellent comparative model for studying the development and function of blood vasculature involved in gas exchange *in vivo.* We performed scRNAseq on isolated adult zebrafish gills and identified a variety of cell clusters corresponding to cell types found in mammalian lungs.

Zebrafish gill epithelial cells share key genes with similar epithelial cell types found in mammalian lungs. Our single-cell analysis revealed rhcga+, col4a4+, and notch3+ (stalk) epithelial clusters, as well as muc13a+ (mucosal) epithelial clusters (**Fig. 3A-F**). The Rh family C glycoprotein (RHCG) is expressed in lung bronchial epithelial cells, contributing to ammonia metabolism and transport (Han et al., 2009), while lung alveolar epithelial cells express type IV collagen (Simon et al., 1993). Gill and lung type IV collagen-producing epithelial cells are both tightly associated with gas-exchange capillary cells, forming the critical gas-exchange blood-gas barrier (Loscertales et al., 2016). Notch3+ gill epithelial cells resemble notch+ lung suprabasal cells, which play an important role in fate determination during development and repair following lung injury (Bodas et al., 2022; Deprez et al., 2020; Hewitt & Lloyd, 2021; Mori et al., 2015). Murine MUC13 is a specific marker of lung pneumocytes and bronchiolar epithelium (Williams et al., 2001), similar to zebrafish gill muc13a+ mucosal epithelial cells. These and other similarities suggest that lungs and gills share conserved epithelial features.

We also found similarities between other specialized cell types in gills and lungs. Mucus-producing MUC5/*muc5.1*-positive goblet cells are found in both lungs and gills, respectively (**Fig. 3G,H**), and both express SPDEF, a transcription factor necessary and sufficient for mammalian goblet cell differentiation (Rajavelu et al., 2015). Gill SCC cells (**Fig. 3L,M**) specifically express transcription factor *pou2f3* (**Fig. 3B**), the “master regulator” of mammalian tuft cell gene expression and development (Wu et al., 2022), suggesting differentiation of these specialized mucosal chemosensory cells may also be conserved between the two gas-exchange organs. Ionocyte subtypes in gills and lungs also share the expression of comparable key ion transporter genes (**Fig. 3A,B,J,K**). Mammalian type A ionocytes express the voltage-gated chloride channel CLCNKA (Yuan et al., 2023) whereas zebrafish KS ionocytes express a very similar voltage-gated *chloride channel 2c (clcn2c*). Mammalian type B pulmonary ionocytes express numerous proton pump genes (ATP6AP2 and ATP6V1F) while gill HR ionocytes express orthologs of many of these same proton-pumping genes, including *atp6v1a* (Kowalewski et al., 2021). Gill oxygen-sensing neuroepithelial cells share expression of sensory signaling molecules such as serotonin (**Supp. Fig. 3E,F**), as well as oxygen-sensing function and endodermal tissue origins, with mammalian pulmonary neuroendocrine cells (Candeli & Dayton, 2024; Hockman et al., 2017; Jonz et al., 2004; Pan et al., 2022). Together, these examples suggest that lungs and gills may be more similar in terms of their cellular organization and function than previously recognized.

The two gas-exchange organs also display parallels in their vasculature and endothelial cell composition. Like lungs, gills have both lymphatic and blood vascular networks. Lymphatic vessels are only found adjacent to the cartilaginous core inside the gill filament stalks (**Fig. 6**). Gill lymphatics specifically express the “diagnostic” *mrc1a, flt4, and lyve1b* genes and their corresponding *Tg(mrc1a:eGFP)^y251^* (Jung et al. 2017)*, Tg(5.2lyve1b:dsred)^nz101^* (Okuda et al., 2012), and *Tg(flt4:yfp)^hu4881^* (Hogan et al., 2009) transgenic reporters. Each gill filament stalk also contains two blood vessels, an afferent filamental artery (deoxygenated, flowing in a proximal to distal direction through the stalk) and an efferent filamental artery (oxygenated, flowing in a distal to proximal direction through the stalk) (**Fig. 4B,C**). Blood exits afferent filamental arteries into each of the lamellae, returning to the circulation via efferent filamental arteries. The filamental arteries and lamellar rim, but not the center of the lamellae, are marked by arterial *Tg(kdrl:mcherry)^y206^* transgene expression. Single-cell analysis reveals distinct molecular identities within different parts of the gill arterial endothelium, as evidenced by differential expression of the *angpt2a, ihha, aqp8a.1,* and *vwf* genes (**Fig. 5D-H**). At this point, it remains unclear, however, whether the different genes expressed in different parts of the gill arterial endothelium reflect fundamental distinctions in the underlying differentiation of their resident endothelial cells, or altered expression due to differences in their local environments (e.g., blood oxygenation, type of flow).

In addition to arterial and lymphatic endothelial cells, the largest single endothelial cell cluster identified in our zebrafish gill single-cell analysis, by far, is a capillary cluster with a gene expression profile resembling the Cap2 or “Aerocyte” endothelial cells found in mammalian lung alveolae (**Fig. 7A,B, Supp. Fig. 7A**) (Ellis et al., 2020; Gillich et al., 2020; Niethamer et al., 2020). HCR *in situ* hybridization for *ncam3,* the zebrafish ortholog of mammalian marker Ncam1, reveals that these gill zAerocytes represent the endothelial cells of the gas-exchange lamellae, with the exception of the lamellar rim noted above (**Fig. 7C-E**). Previous studies have noted these same lamellar cells, calling them “pillar cells” due to their unique structure (Evans et al., 2005; Wilson & Laurent, 2002), though detailed molecular and volume ultrastructural characterization of these cells has not been carried out. Our FIB-SEM studies, coupled with extensive segmentation, show that the *ncam3*+ zAerocytes are deeply penetrated by cellular protrusions from adjacent epithelial cells (**Fig. 8Q, Supp. Fig. 10**). The presence of intracellular actin- and myosin-enriched assemblies surrounding the pillars suggests that they may have a contractile function (**Fig. 8K-M, Supp. Fig. 8**) that could potentially contribute to the structural and/or physiological adaptations required for effective gas-exchange under different environmental conditions, for example, reduced oxygen (hypoxia).

In the study by Gillich et al. (2020), the authors examined gas-exchange capillary cells from ancestral amniote species—the American alligator (Alligator mississippiensis) and Western painted turtle (Chrysemys picta bellii)—and found that the gas-exchange capillary cells of both species co-expressed markers from mammalian lung Aerocytes (Cap2) and general capillary cells (Cap1). However, in the zebrafish gill, zAerocytes did not co-express markers of both lung Aerocytes (Cap2) and general capillary cells (Cap1). Instead, in zebrafish gills, zAerocytes strongly express markers of lung Aerocytes (Cap2), while other gill cells express lung general capillary (Cap1, gCap) cell markers (**Supp. Fig. 7A**).

A small additional “hybrid capillary-epithelial” cluster was also identified in our single-cell analysis (**Supp. Fig. 11**). Given the very tight association of zAerocytes with, and particularly their deep penetration by, adjacent lamellar epithelial cells, we speculate that this cluster is the result of doublet formation between these cells due to shearing off of epithelial protrusions into zAerocytes during single-cell dissociation. Indeed, the epithelial-specific transcripts found in the “hybrid” cells, including *col4a3, col4a4 (type IV collagen), ca6 (carbonic anhydrase), sema3fa (semaphorin 3), and epcam,* are all highly expressed in the *col4a4*+ epithelial cells located immediately adjacent to zAerocytes in gill lamellae (**Figs. 3B,P, 5B-C**). Furthermore, a “computational mixing experiment” artificially generating a doublet cell cluster containing gene transcripts from both zAerocyte and *col4a4*+ epithelial cell clusters shows that the hybrid cell cluster strongly resembles the computationally-generated doublets and does not contain uniquely enriched transcripts of its own, strongly supporting the idea that these transitional “hybrid” cells are the result of zAerocyte/epithelial cross-contamination (**Supp. Fig. 11**). The appearance of “doublets” due to incomplete dissociation of tightly adhering cell populations is a known concern when performing single-cell RNA sequencing experiments, and our findings suggest that results should be interpreted with caution and additional rigorous methods may be needed in order to definitively assess the molecular characteristics of the very closely apposed endothelial and epithelial cell populations present in gas exchange organs.

Together, our study and previous reports suggest that gills and lungs share a wide range of specialized cell types, including epithelial, mucosal, chemosensory, and endothelial cells. Many of these cells share key transcription factors and/or genes that are critical for their shared functional properties. The conserved molecular characteristics of arteries, lymphatics, and capillaries in the gills and lungs suggest that studying the vasculature in the accessible gill model may provide important new insights into the origins and function of lung endothelial cell populations. Zebrafish tissues and organs are also highly regenerative, including the gills (Cadiz et al., 2024; Jonz, 2024), making this a potentially powerful model for investigating mechanisms that promote endothelial regeneration after injury.

## Study Approval

Zebrafish husbandry and research protocols were reviewed and approved by the NICHD Animal Care and Use Committee at the National Institutes of Health. All animal studies were conducted following NIH-approved protocols and in compliance with the Guide for the Care and Use of Laboratory Animals.

## Data and Materials Availability

Raw and processed 10X scRNA-seq data can be found on GEO using accession number GSE#. The code used to analyze and visualize the data can be found at: https://github.com/nichd-Weinstein/. All other data reported in this manuscript are available from the corresponding author upon request.

## Acknowledgments

The authors would like to thank members of the Weinstein laboratory for their critical comments on this manuscript. The Authors would also like to thank the Research Animal Branch of the *Eunice Kennedy Shriver* National Institute of Child Health and Human Development and the RAMB contract animal management staff for excellent animal care and husbandry. This research was supported by the Intramural Research Program of the *Eunice Kennedy Shriver* National Institute of Child Health and Human Development (ZIA-HD008915, ZIA-HD008808, and ZIA-HD001011, to BMW, and ZIC-HD008986 to RKD) and by the Intramural Research Program of the National Cancer Institute (to KN).

