## Supplemental Data for "Specialized gas-exchange endothelium of the zebrafish gill"

#### SUPPLEMENTAL FIGURE LEGENDS

**Supplemental Figure 1. Gill filament histology.**

**A,B,** Brightfield micrograph of an adult zebrafish gill filament longitudinal section stained with Alcian Blue, revealing gill cartilage in the core of the filament in blue. Panel B shows a higher magnification of the boxed area in panel A. **C,D,** Brightfield micrograph of an adult zebrafish gill filament longitudinal section stained with H&E, revealing gill cell nuclei in dark purple and cytoplasm/extracellular matrix in pink. Panel D shows a higher magnification of the boxed area in panel C. **E,F,** Brightfield micrograph of an adult zebrafish gill filament longitudinal section with Movat pentachrome, revealing cartilage in the core of the filament in blue-green, tissue surrounding cartilage in yellow, and lamellar tissue in brown. Panel F shows a higher magnification of the boxed area in panel E. Scale bars = 150 µm (A,C,E), 25 µm (B,D,F).

**Supplemental Figure 2. Gill-associated cell clusters and their identities.**

**A,** UMAP of 29 zebrafish gill-associated cell clusters and their identities. **B,** Table of cell identities and their identifying genes.

**Supplemental Figure 3. Gill epithelial cell types.**

**A,B,** (A) Confocal micrograph of a transverse section of an adult zebrafish gill filament subjected to hybridization chain reaction (HCR) for *clcn2c* (green), *rhcgb* (magenta), and *trpv5* (orange), revealing *en face* gill lamellae, and (B) higher magnification view of the boxed area in panel A. **C,D,** (C) Confocal micrograph of a transverse sections of adult zebrafish gill filament subjected to HCR for *adgrf3a* (green) and *aldh1a3* (magenta), revealing *en face* gill lamella, and (D) higher magnification view of the boxed area in panel C. **E,F,** (E) Lateral view confocal micrograph of adult zebrafish gill filaments subjected to immunofluorescence for oxygen-sensing neuroepithelial cells (anti-serotonin Ab) and neuronal cells (anti-zn-12), and (F) higher magnification view of the boxed area in panel E. Scale bars: 50 µm (A,C), 25 µm (B,D,F), 100 µm (E).

**Supplemental Figure 4. Gill filament array tomography.**

**A-D,** (A) Array tomography micrograph slice showing *en face* gill lamella view with erythrocytes in the afferent (deoxygenated blood entering the gill lamella; yellow box) and efferent (oxygenated blood exiting the gill lamella; blue and red boxes) filamental arteries. Boxes note areas shown in higher magnification in panels B-D. (B) higher magnification image of the efferent filamental artery (asterisk) taken from a different plane in the array tomography dataset than panel A. (C) higher magnification image of the efferent filamental artery (asterisk) showing lateral branches (arrows) into the lamellar leaflets. (D) higher magnification image of the afferent filamental artery (asterisk) with a lateral branches (arrows) into the lamellar leaflet. **E,F,** (E) Array tomography micrograph slice showing *en face* gill lamella view with erythrocytes in the gas-exchange intralamellar space, and (F) higher magnification image of the boxed area in panel E showing erythrocytes (arrow) threading their way through the lumen between cells with large irregular nuclei (red asterisk). Scale bars = 10 µm (A-F).

**Supplemental Figure 5. Confocal imaging of gills with endothelial-specific transgenics.**

**A,B,** Confocal micrographs of transverse sections of adult *Tg(kdrl:mcherry)*$^{y206}$ transgenic zebrafish gill filaments showing *en face* gill lamella for (A) kdrl:mcherry only or (B) kdrl:mcherry and Hoechst-labeled erythrocytes (zebrafish erythrocytes are nucleated). EFA and AFA denote efferent and afferent filamental arteries, respectively. Small arrows indicate kdrl+ lamellar rim, asterisk notes kdrl- center of lamellar leaflet. **C-H,** Confocal micrographs of vessels (green) in adult *Tg(fli1a:eGFP)*$^{y1}$ transgenic zebrafish gill filaments, showing lateral (C,D), dorsal (E), and transverse section (F-H) views. Panel D is the same as E but includes erythrocyte autofluorescence. Asterisks in panels C and F note fli1a:egfp-low central areas of the lamellae. Panels G and H show fli1a:egfp expression in immature distal lamellar leaflets (H shows higher magnification image of the boxed region in G). Scale bars = 25 µm (A,F,H), 50 µm (B,C,D,E,G).

**Supplemental Figure 6. Confocal imaging and array tomography of gill lymphatics**

**A,B,** Confocal micrographs of arterial (magenta) and lymphatic (green) endothelium in transverse sections of adult *Tg(kdrl:mcherry)*$^{y206}$, *Tg(mrc1a:eGFP)*$^{y251}$ double transgenic zebrafish gill filaments, revealing *en face* gill lamellae. **C-E,** (C) Array tomography micrograph showing a transverse section through a gill filament stalk with lymphatic vessels surrounding the cartilaginous core, (D,E) higher magnification image of the boxed area in C, without (D) or with (E) lymphatic vessels pseudocolored green. **F-H,** (F) Array tomography micrograph slice showing a transverse section through a gill filament stalk with a white blood cell present inside the lymphatic vessel, (G,H) higher magnification image of the boxed area in F, without (G) or with (H) the lymphatic vessel and white blood cell pseudocolored in green and red, respectively. Scale bars = 50 µm (A,B), 10 µm (C-H).

**Supplemental Figure 7. Gill zAerocytes**

**A,** Dot plots comparing gene expression in zebrafish gill *ncam3+* ECs to other gill ECs, and comparing gene expression in mouse lung aerocyte (Cap2), general capillary (Cap1, gCap), and other lung ECs. Gill ncam3+ aerocytes express Cap2 but not Cap1 markers. **B-F,** Lateral view confocal micrographs of adult *Tg(fli1a:eGFP)*$^{y1}$ transgenic zebrafish gill filaments subjected to HCR for ncam3. Images are from the same field, and they show endothelium (A-C, green), autofluorescent erythrocytes (B,C,E, blue), and/or ncam3 staining (C,D, magenta). Partially folded gill lamellae revealing the fli1a:egfp low, ncam3-positive central gas-exchange area of the lamellae. Scale bars = 100 µm (A-E).

**Supplemental Figure 8. Confocal imaging of gill zAerocyte cellular structures**

**A-F,** Confocal micrographs of adult *Tg(fli1a:eGFP)$^{y1}$* transgenic gill lamellar zAerocytes (green) with Hoechst labeled nuclei (blue), showing zAerocytes in the tip of a lamellar leaflet (A,C,E) and higher magnification images of a single zAerocyte (B,D,F) from the boxed region in panel A. Panels include flia:egfp and Hoechst (A,B), flia:egfp only (C,D), or Hoechst only (E,F) images. **G-O,** Confocal micrographs of actin-rich structures (green) in gill lamellar zAerocytes in *Tg(fli1a:lifeact-egfp)* transgenic adults injected intravascularly with BODIPY546 nuclear membrane-labeling dye (magenta). Images show a single optical section of zAerocytes in the tip of a lamellar leaflet (G,J,M), higher magnification images of a single zAerocyte (H,K,N) from the boxed region in panel G, and lateral view 3D reconstructions (I,L,O) the confocal data for the same zAerocyte shown in panel H. Panels include fli1a:lifeact-egfp and BODIPY546 (G,H,I), fli1a:lifeact-egfp only (J,K,L), or BODIPY546 only (M,N,O) images. **P-X,** Stimulated emission depletion (STED) micrographs of actin-rich structures (green) and nuclei (blue) in gill lamellar zAerocytes stained with Abberior's STED STAR-dye conjugated phalloidin and Live DNA dye, respectively. Images show dorsal views of a zAerocyte and surrounding erythrocytes (P,S,V), a higher magnification dorsal view of the same zAerocyte (Q,T,W), and lateral views of one half of the same zAerocyte (R,U,X). Panels include phalloidin and Hoechst(G,H,I), phalloidin only (J,K,L), or Hoechst only (M,N,O) images. **Y,Z,** Confocal micrographs of anti-phosphorylated myosin light chain immunofluorescence (Phospho-MLC; magenta) and erythrocyte autofluorescence (RBC autofluorescence; white) in adult zebrafish gill filaments. Panels show (Y) an overview image of gill filaments and lamellae, and (Z) a higher magnification phospho-MLC only image of the boxed area in Y, revealing enrichment of phospho-MLC in zAerocyte pillars. Scale bars = 5 µm (A,C,E,G,J,M), 2 µm (B,D,F,H,I,K,L,N,O,P-X), 10 µm (Z), 50 µm (Y).

**Supplemental Figure 9. FIB-SEM imaging of zAerocyte "pillars"**

**A-C,** FIB-SEM 3D reconstruction of a single gill zAerocyte from (A) dorsal view with pillar openings circled in blue, (B) lateral view from outside of cell, (C) view of pillars from the inside of the cell (area noted by box in panel B). Scale bar = 1 µm (B).

**Supplemental Figure 10. FIB-SEM imaging of epithelial penetration into zAerocytes**

**A-C,** FIB-SEM 3D reconstruction of a single gill zAerocyte from (A) lateral view, (B) dorsal view, and (C) higher magnification of the boxed area in panel B. Red lines and arrows in panel B indicate slices shown in either panels D-F, G-I, or J-L. **D-L,** (D,G,J) FIB-SEM sections corresponding to the red lines in panel B and red arrows in panel C, (E,H,K) the same FIB-SEM sections with zAerocytes pseudocolored in green and epithelial cells pseudocolored in blue, (F,I,L) higher magnification images of the boxed areas in panel E, H, and K, respectively. Blue arrows in panels F, I, and L indicate epithelial protrusions into zAerocyte "pillars", while red arrows indicate areas within pillars containing only extracellular matrix. Scale bars = 1 µm (A,D-L).

**Supplemental Figure 11. Presence of cells with hybrid zAerocyte-epithelial identity**

**A,B,** (A) UMAP projection of all zebrafish gill-associated cells, and (B) magnified view of selected cell clusters boxed in red in panel A, showing *col4a4*+ epithelial cells in orange, zAerocytes in blue, and cells displaying a hybrid identity in pink. **C,** UMAP projection of sub-clustered endothelial cells (ECs), showing the same hybrid cell cluster adjacent to the zAerocyte cluster. **D,** Computational cluster similarity comparison between zAerocyte-epithelial "hybrid" cells and (1) the same hybrid cell cluster, (2) simulated zAerocyte/col4a4+ epithelial doublets, (3) zAerocytes, (4) col4a4+ epithelial cells, or (5) randomly selected non-aerocyte/col4a4+ epithelial/hybrid cells from the entire scRNAseq data set, where distance is the Euclidean distance between cells in gene expression space. See "Hybrid cell doublet analysis" in the Materials and Methods section for additional details.

#### SUPPLEMENTAL MOVIES

**Supplemental Movie 1**

Overview of gill structures and blood flow in adult Tg(kdrl:eGFP) (arteries) and Tg(gata1:dsRed) (blood cells) zebrafish.

**Supplemental Movie 2.**

A close-up *en face* view of the gill lamellae in an adult Tg(kdrl:mCherry) (artery) zebrafish injected with Hoechst (nuclear) dye.

**Supplemental Movie 3.**

Array tomography segmentation and 3-D reconstruction of the blood endothelium luminal spaces, erythrocytes, and epithelial layers.

**Supplemental Movie 4.**

A close-up 3D rendering of adult Tg(kdrl:mCherry) (artery) and Tg(mrc1a:eGFP) zebrafish gill filaments.

**Supplemental Movie 5.**

Pseudocolored array tomography sections revealing afferent (purple) and efferent arteries (purple) cross-sections, cartilage core (blue), lymphatic vessels (green) surrounding the cartilage, and a white blood cell (pink) within the lymphatic vessel.

**Supplemental Movie 6.**

A close-up 3D rendering of adult Tg(fli1:eGFP) (pan-endothelial) zebrafish gill filament treated with HCR for *ncam3*, a gill zAerocyte marker, and blue autofluorescence that marks red blood cells.

**Supplemental Movie 7.**

Gill lamellae gas-exchange endothelium live imaging – showing DRAQ5 nuclear stain, revealing pseudocolored green zAerocytes and magenta red blood cells.

**Supplemental Movie 8.**

A close-up 3D rendering of adult Tg(fli1:LifeAct-eGFP) zebrafish gill lamellae labeled with BODIPY546 membrane dye.

**Supplemental Movie 9.**

A 3D rendering of FIB-SEM segmented zebrafish gill zAerocyte, revealing unique morphological structures of the gas-exchange cell.

**Supplemental Movie 10.**

A 3D rendering of array tomography segmented zebrafish gill lamella, showing red blood cells within the afferent and efferent filamental arteries and capillary lumens, as well as Aerocyte and epithelial cell nuclei within the lamella.

#### SUPPLEMENTAL FILES

##### Supplemental File 1.

Complete ultrastructural reconstruction of an individual zAerocyte using FIB-SEM, revealing "Wheel-hub-like" shaped cell.

#### Supp. Fig. 1. Gill filament histology

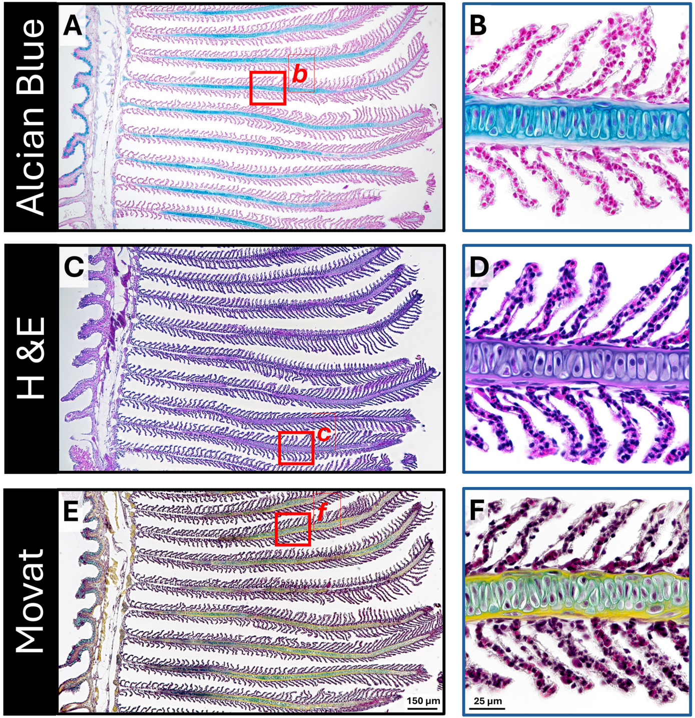

**Supp. Fig. 2. Gill-associated cell clusters and their identities**

**A**

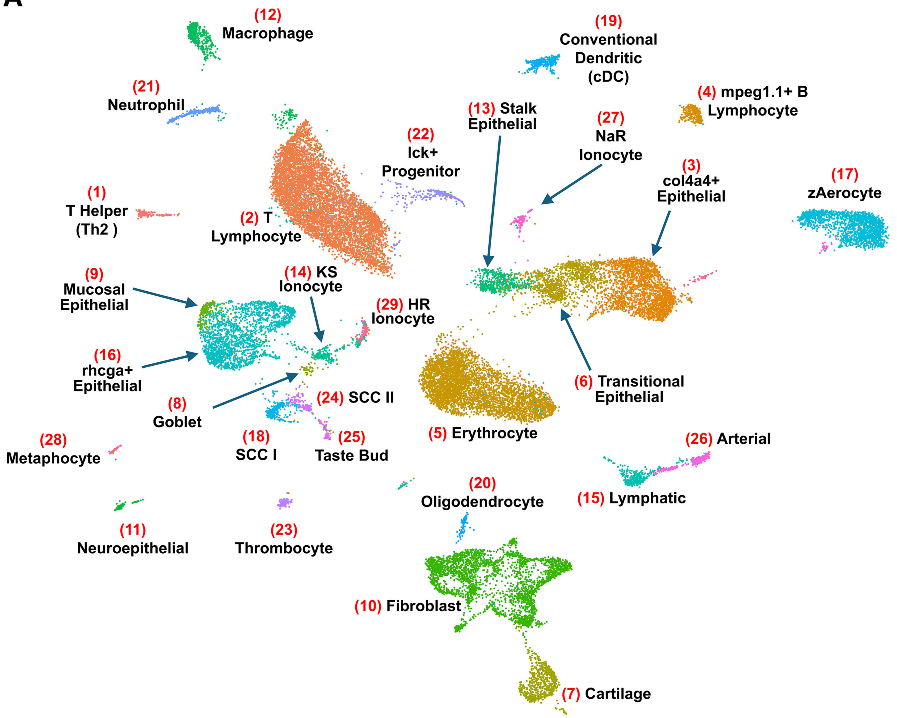


**B**

| | IDENTITY | IDENTIFYING GENES | | | |
|---|---|---|---|---|---|
| | | 1 | 2 | 3 | 4 |
| 1 | T helper (Th2) | sox13 | il13 | il1rl1 | il4 |
| 2 | T Lymphocyte | ccr7 | zap70 | cd8a | foxp3a |
| 3 | col4a4+ Epithelial | lamb2l | col4a4 | kcp | col4a3 |
| 4 | mpeg1.1+ B Lymphocyte | igic1s1 | igl1c3 | igl3v5 | ighv1-4 |
| 5 | Erythrocyte | creg1 | alas2 | cahz | hemgn |
| 6 | Transitional Epithelial | apoeb | fat2 | cldni | egfra |
| 7 | Cartilage | acana | col2a1a | col11a2 | col11a1a |
| 8 | Goblet | muc5.1 | foxa3 | SPDEF | agr2 |
| 9 | Mucosal Epithelial | lye | scel | muc13a | evplb |
| 10 | Fibroblast | lum | col1a2 | col5a1 | col1a1a |
| 11 | Neuroepithelial | slc18a2 | tph1a | sox2 | sox1b |
| 12 | Macrophage | mfap4 | c1qb | c1qc | cx32.2 |
| 13 | Stalk Epithelial | cldna | tnfrsf9a | notch3 | grhl2a |
| 14 | KS Ionocyte | slc12a10.2 | kcnj1a.5 | kcnj1a.6 | atp1a1a.2 |
| 15 | Lymphatic | flt4 | mrc1a | lyve1b | cxcl12a |
| 16 | rhcga+ Epithelial | rhcga | ponzr3 | pkhd1l1 | krt98 |
| 17 | zAerocyte | pecam1b | ncam3 | itga1 | acvrl1 |
| 18 | Solitary Chemosensory (SCC I) | igsf9a | alox5a | ltc4s | adgrg11 |
| 19 | Conventional Dendritic (cDC) | ccl35.1 | cd83 | xcr1a.1 | ccl35.2 |
| 20 | Oligodendrocyte | mbpa | mpz | plp1b | sox10 |
| 21 | Neutrophil | mpx | il6r | cfbl | mmp9 |
| 22 | lck+ Progenitor | mki67 | cdk1 | top2a | stmn1a |
| 23 | Thrombocyte | itga2b | mpl | thbs1b | vegfc |
| 24 | Solitary Chemosensory (SCC II) | aldh1a3 | tuba1a | vil1 | SH2D6 |
| 25 | Taste Bud | ano2 | vill | gng13a | scg3 |
| 26 | Arterial | vwf | aqp8a.1 | angpt2a | efnb2b |
| 27 | NaR Ionocyte | trpv6 | s100a11 | igfbp5a | gcm2 |
| 28 | Metaphocyte | grn2 | spic | il22ra2 | cd180 |
| 29 | HR Ionocyte | rhcgb | ceacam1 | slc9a3.2 | slc4a1b |

#### Supp. Fig. 3. Gill epithelial cell types

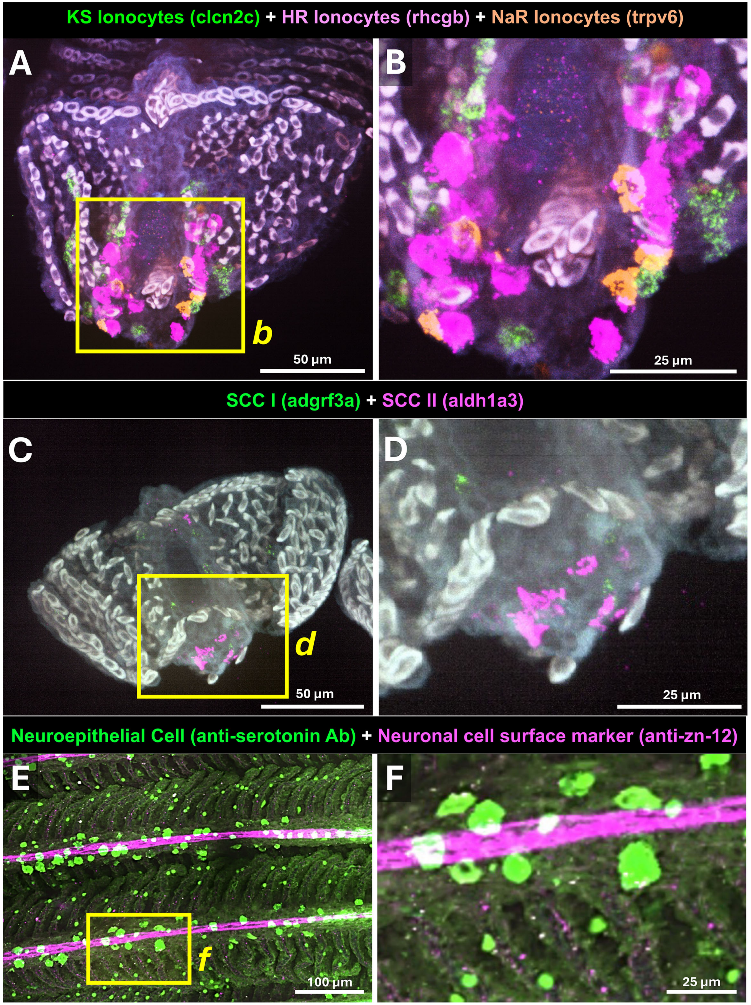

**Supp. Fig. 4. Gill filament array tomography**

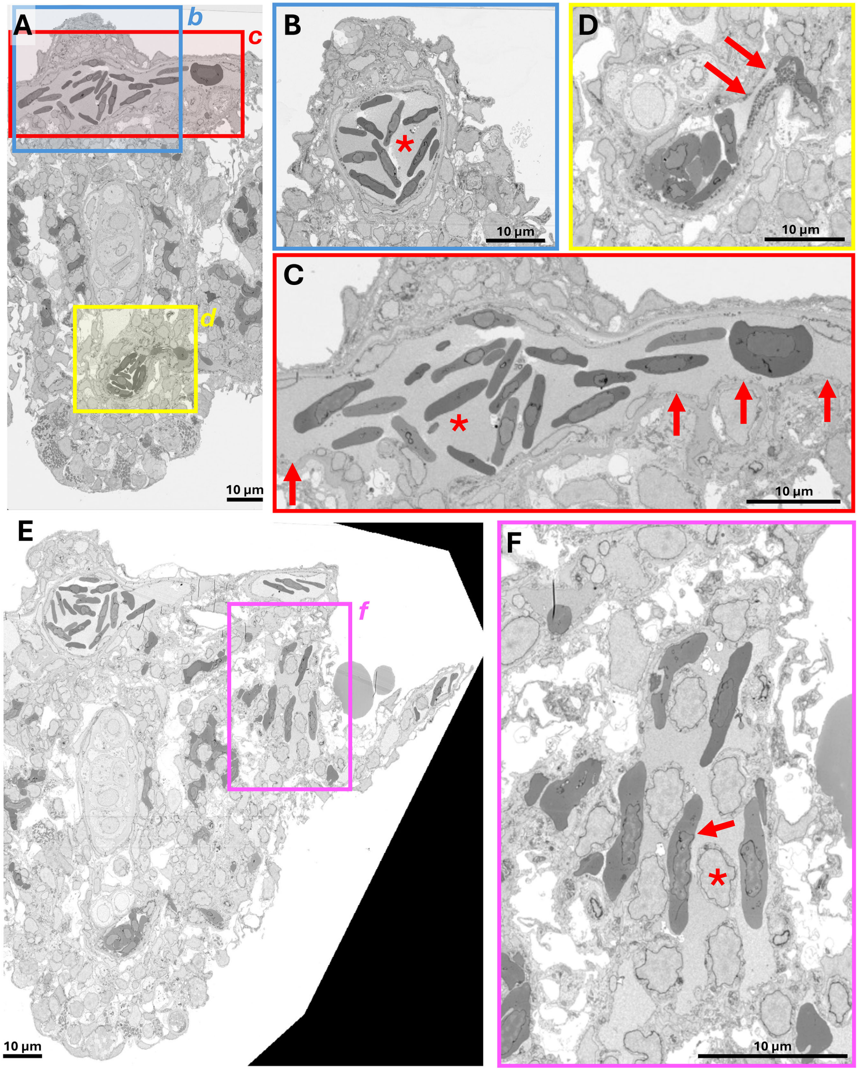

### Supp. Fig. 5. Confocal imaging of gills with endothelial-specific transgenics

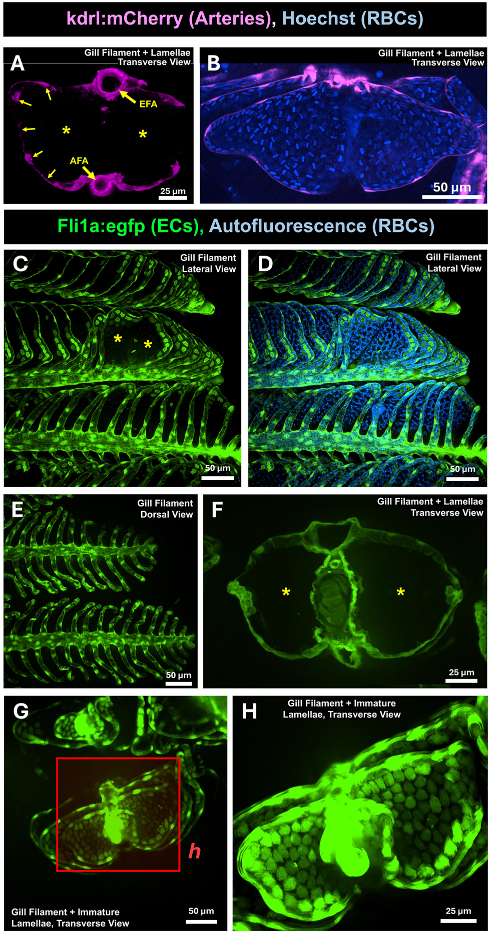

#### Supp. Fig. 6. Confocal imaging and array tomography of gill lymphatics

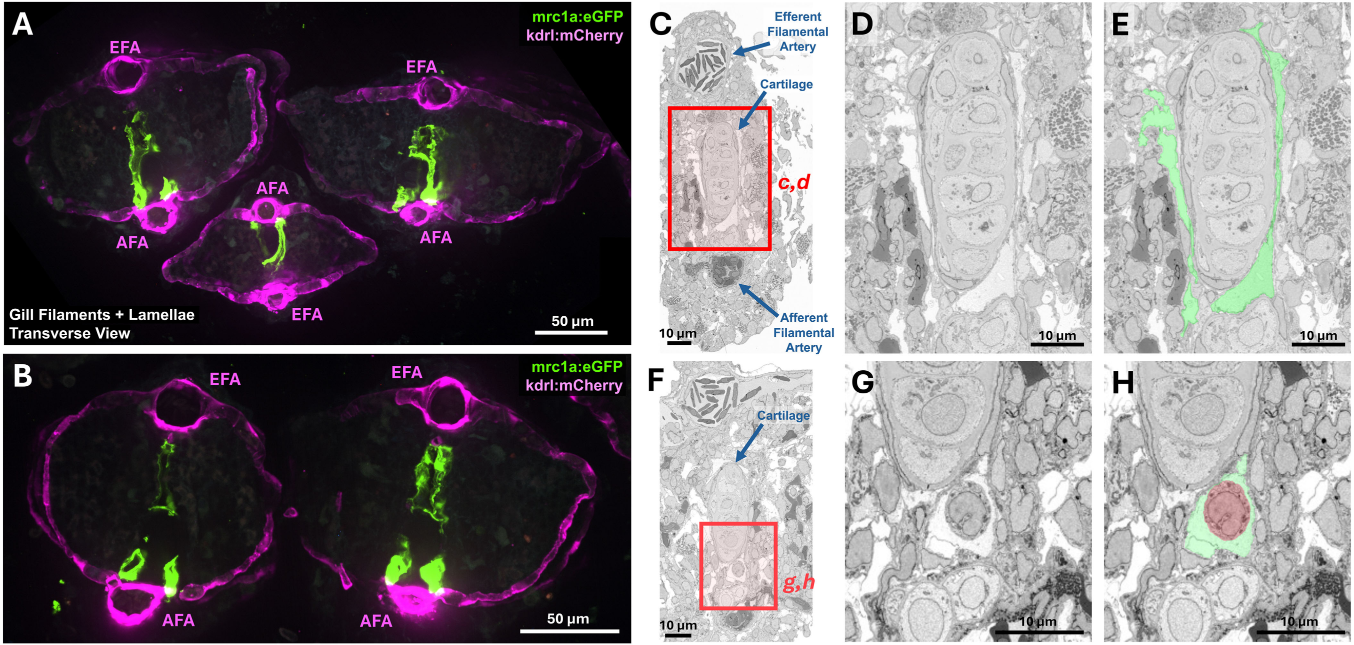

#### Supp. Fig. 7. Gill zAerocytes

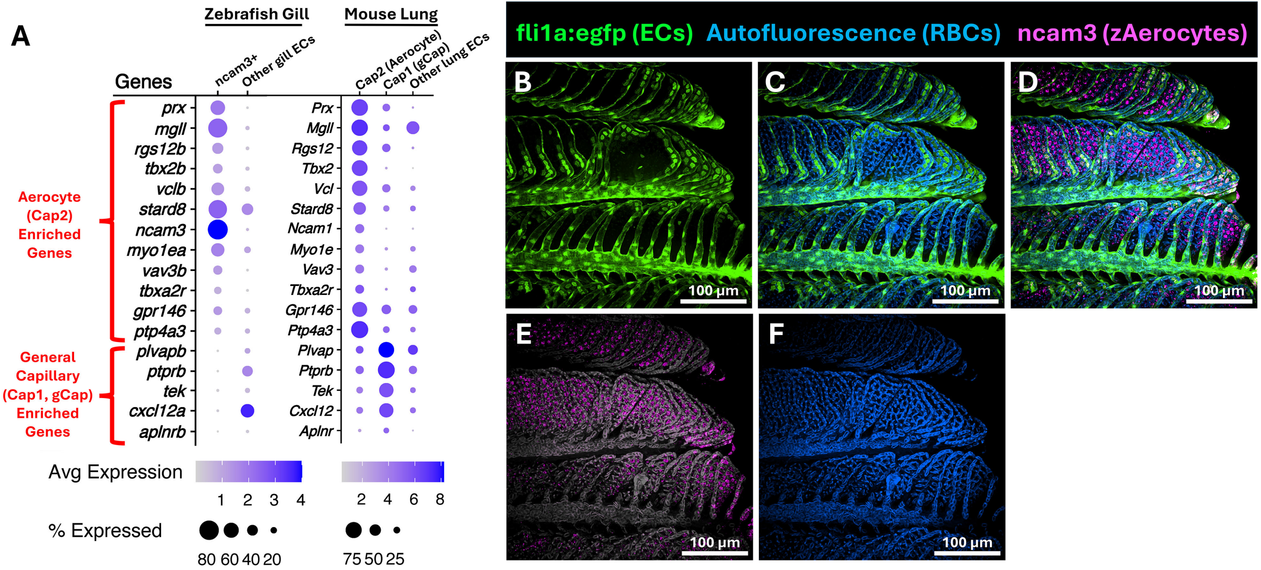

**Supp. Fig. 8. Confocal imaging of gill zAerocytes**

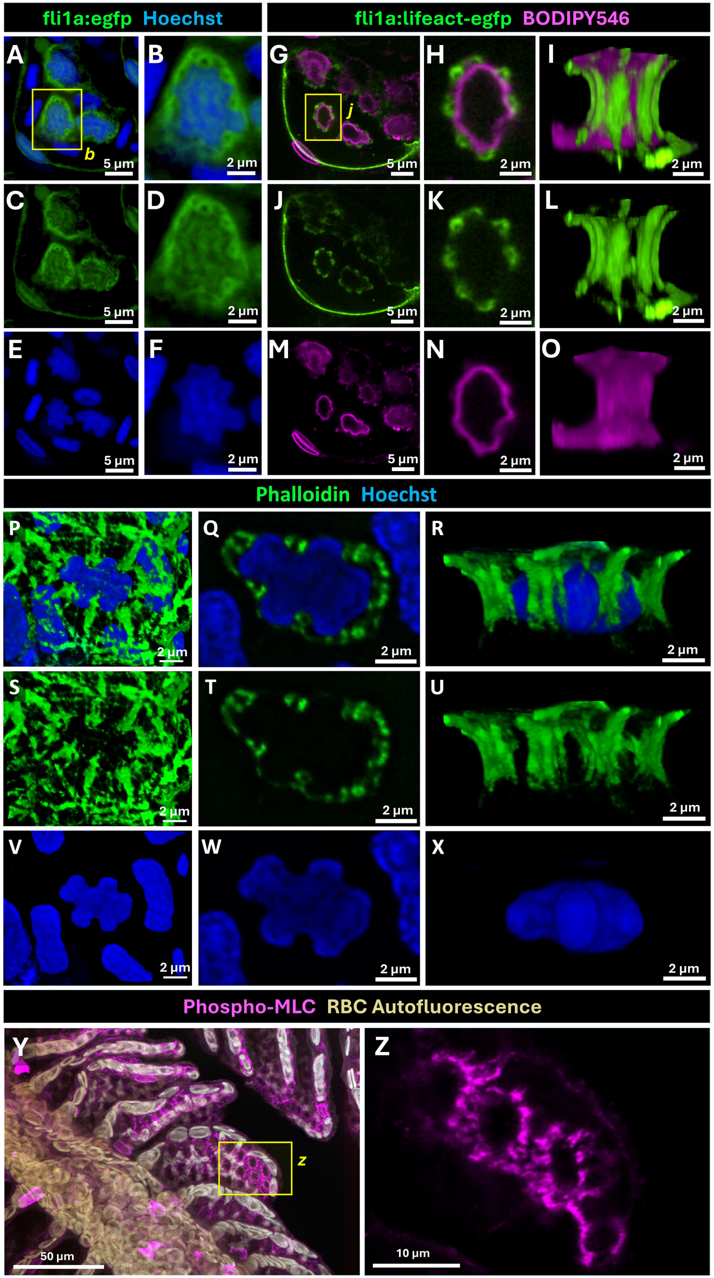

### Supp. Fig. 9. FIB-SEM imaging of zAerocyte "pillars"

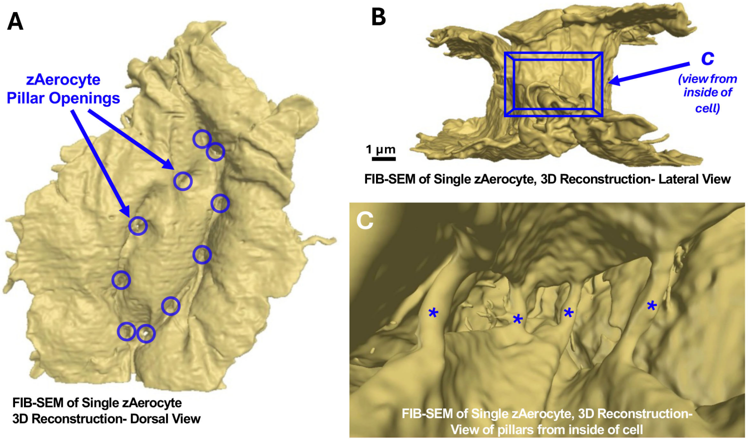


FIB-SEM of Single zAerocyte
3D Reconstruction- Dorsal View

FIB-SEM of Single zAerocyte, 3D Reconstruction- Lateral View

FIB-SEM of Single zAerocyte, 3D Reconstruction-
View of pillars from inside of cell

**Supp. Fig. 10. FIB-SEM imaging of epithelial penetration into zAerocytes**

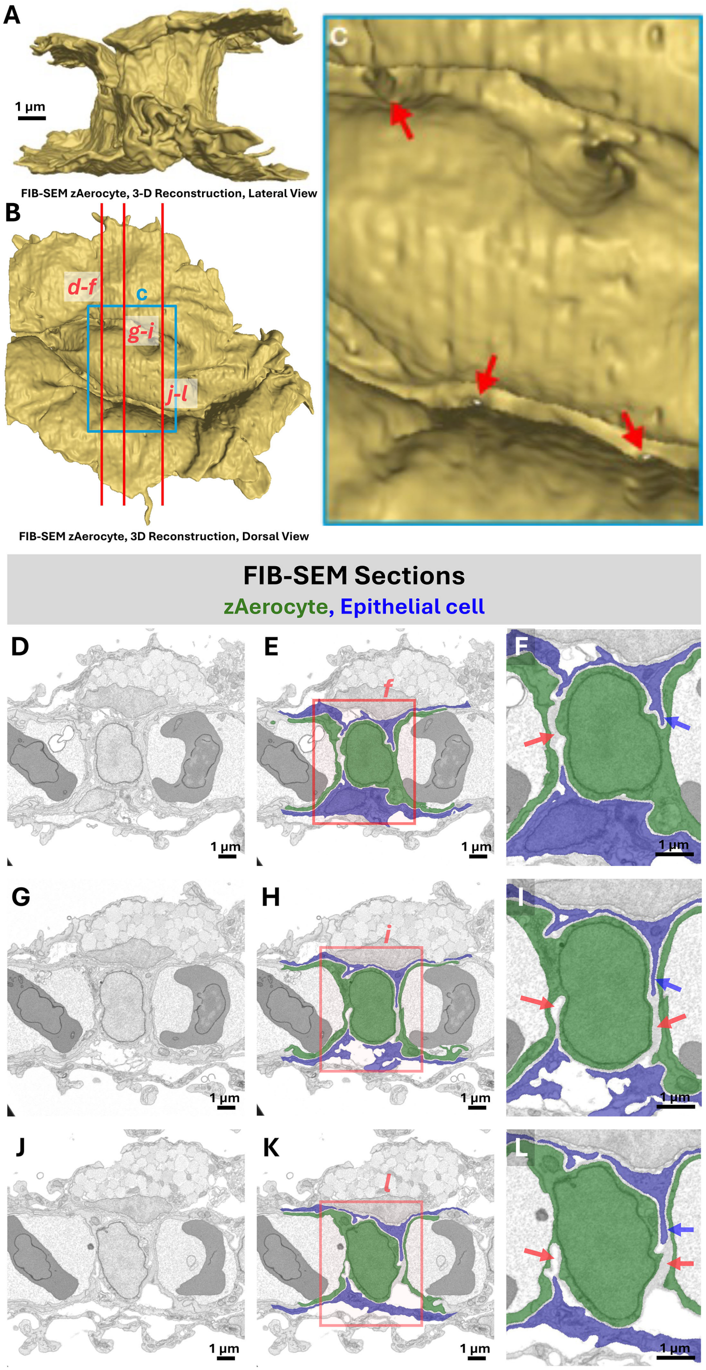

### Supp. Fig. 11. Presence of cells with hybrid zAerocyte-epithelial identity

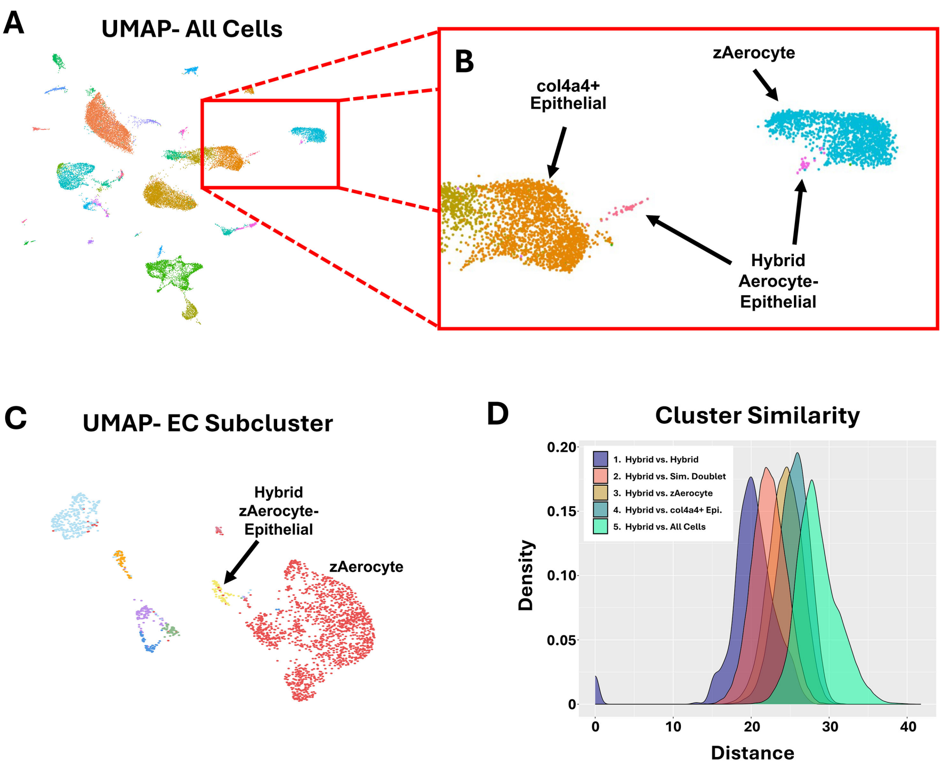
